# Ybx1 Deficiency Causes ROS-Driven IBD-Like Intestinal Inflammation and Postnatal Lethality

**DOI:** 10.64898/2026.01.26.701719

**Authors:** Bo Zhu, Lakhansing Pardeshi, Yingying Chen, Xianqing Zhou, Wei Ge

## Abstract

Y box-binding protein 1 (YB-1; Ybx1/*ybx1*) is essential for zebrafish development. Maternal *ybx1−/−* mutants exhibited embryonic lethality, whereas zygotic mutants (Z*ybx1*−/−) showed high postnatal mortality between 10 and 20 dpf, although a small fraction survived to adulthood. Western blot and immunohistochemical analysis revealed strong, transient expression of Ybx1 in intestinal enterocytes from 3 to 5 days post-fertilization (dpf), followed by rapid ubiquitin-mediated degradation at 6 dpf, coinciding with defective intestinal development and compromised gut homeostasis. RNA-seq analysis identified elevated reactive oxygen species (ROS) and upregulation of matrix metalloproteinases mmp9 and mmp13a in Z*ybx1*−/− larvae. Antioxidant treatment with ascorbic acid rescued postnatal lethality and alleviated intestinal defects, whereas prooxidant exposure exacerbated them. Pharmacological inhibition of Mmp9 or Mmp13a similarly prevented lethality, highlighting a ROS–MMP axis driving tissue damage. By 30 dpf, surviving mutants exhibited progressive intestinal impairment and severe pathology. These findings demonstrate that Ybx1 deficiency triggers ROS-dependent intestinal inflammation, MMP-mediated gut damage, and postnatal lethality, establishing Ybx1-deficient zebrafish as a robust model for studying inflammatory bowel disease (IBD)-like intestinal disorders.

## Introduction

The Y-box binding protein (YBP) family, a subset of the evolutionarily conserved cold shock protein (CSP) superfamily, plays critical roles in various cellular processes, including transcription, translation, cell proliferation, and DNA repair ^1–3^. In vertebrates, the YBP family typically includes three members: YB-1, YB-2, and YB-3. In mice, these are designated MSY1 (YB-1), MSY2 (YB-2), and MSY4 (YB-3) ^4^. MSY1-deficient mice exhibit embryonic lethality; nonetheless, the mutant embryos (*YB-1*−/−) can develop normally up to embryonic day 13.5 (E13.5), but demonstrate lethality between E14.5 and E17.5 ^5,6^. The developmental defects are associated with abnormal patterns of cell proliferation ^5^, likely caused by oxidative stress-induced cell cycle arrest at G1 phase due to increased expression of CDK inhibitors p16 and p21 ^6^. Combined deficiency of MSY1 and MSY4 leads to even earlier embryonic lethality (E8.5 to E11.5) ^7^. In contrast, MSY2-deficient mice are viable but exhibit infertility in both sexes, consistent with its specific expression in germ cells ^8,9^. These findings highlight the essential roles of MSY1 and MSY4 in embryogenesis and MSY2 in fertility.

In contrast to those in vertebrates, invertebrate YBP homologs demonstrate different functional roles. In *Caenorhabditis elegans*, knockout of the YBP homolog (CEYs) causes dysfunction in gametogenesis without lethality ^10^. Similarly, the *Drosophila* YBP homolog Yps is crucial for *oskar* mRNA localization, but *yps*-deficient flies remain viable ^11^. This divergence suggests that while YB-1 is essential for vertebrate embryogenesis, it is not indispensable to development of invertebrates.

Zebrafish possesses a single YBP family member, Ybx1 (*ybx1*), which exhibits structural similarity to mammalian YB-1 but a protein expression pattern similar to mammalian YB-2 ^12^. This suggests that zebrafish Ybx1 may play roles of both YB-1 and YB-2. Previous studies have shown that zebrafish Ybx1 is crucial for maintaining Nodal homeostasis, and its disruption via ENU mutagenesis results in maternal-effect lethality due to gastrulation failure ^13^. Maternal-effect lethality has also been linked to early embryonic defects during the maternal-to-zygotic transition, arising from impaired oocyte maturation ^14^.

Our recent work, using targeted mutagenesis of zebrafish *ybx1*, revealed phenotypes reminiscent of both YB-1 and YB-2 mutants in mammals ^15^. The zygotic *ybx1* mutant (Z*ybx1*−/−) exhibited high post-hatching mortality, with only a small proportion surviving to adulthood, mirroring the embryonic lethality observed in YB-1 mutant mice ^5,6^. Furthermore, surviving female *ybx1* mutants displayed reduced fertility, with ovarian follicle arrested largely at the previtellogenic to vitellogenic transition, analogous to the infertility seen in mouse YB-2 mutants ^8,9^. Subsequent experiments demonstrated that one potential mechanism by which Ybx1 controls early folliculogenesis is to promote follicle cell proliferation by downregulating the cell cycle inhibitor p21 (*cdkn1a*) ^15^.

Following our previous studies on the YB-2-like functions of zebrafish Ybx1 in reproduction or germ cell development, this study aimed to elucidate the functional duality of Ybx1 by examining its YB-1-like activities in development, specifically focusing on the causes of post-hatching mortality in zygotic mutant. We identified a critical period of larval death and, through morphological and biochemical analyses, determined that impaired intestinal development and integrity was likely the primary cause of lethality in *ybx1*−/− mutants. Transcriptomic analysis revealed significantly elevated inflammatory response and proteolysis in the mutant intestine, potentially driven by increased reactive oxygen species (ROS) and elevated expression of the tissue-remodeling enzymes matrix metallopeptidase 9 and 13a (Mmp9/*mmp9* and Mmp13a/*mmp13a*). The phenotypes observed in *ybx1*−/− zebrafish suggest that they may serve as a potential model for human inflammatory bowel disease (IBD).

## Materials and methods

### Zebrafish lines and husbandry

The wild-type (WT) zebrafish (*Danio rerio*) of AB strain and *ybx1*−/− mutants (ZFIN line No.: umo60; generated using TALEN technology) ^15^ were used in this study. The fish were maintained in the ZebTEC Multilinking Rack System (Tecniplast, Buguggiate, Italy) under a controlled condition as follows: water temperature 28 ±1°C, pH 7.5, conductivity 400 *μ*S/cm, and photoperiod of 14-h light:10-h dark. The fish were fed a diet of paramecia, artemia, and artemia+Otohime fish diet (Marubeni Nisshin Feed, Tokyo, Japan) twice daily during the larval, juvenile, and adult stages, respectively. All experiments were conducted under a license from the Government of the Macau Special Administrative Region (SAR) and approved by the Panel on Animal Research Ethics of the University of Macau (Approval No. AEC-13-002).

### Genotyping with high resolution melting analysis (HRMA)

We used HRMA to genotype the zebrafish. The assay was conducted according to the protocol we described previously ^15,16^. A total of about 10,000 zebrafish from crosses of heterozygous parents (+/−) were genotyped. Fish of each genotype (*ybx1*+/+, *ybx1*+/−, and *ybx1*−/−) were subsequently housed in separate tanks for phenotypic analysis.

### Morphological and lethality analysis in larvae

Genotype analysis of the offspring from crosses of heterozygotes (*ybx1*+/−) indicated significant post-hatching lethality in adult *ybx*1−/− mutant. To determine the critical period of lethality, we analyzed fish larvae at four different time points: 5, 10, 15, and 20 days post-fertilization (dpf). Forty larvae were randomly selected at each time point for morphological recording and genotype analysis by HRMA. Prior to imaging, individual larvae were anesthetized with tricaine methanesulfonate (MS-222). The image of each larva was captured using the SMZ18 microscope (Nikon, Tokyo, Japan). Three independent groups of 40 larvae were analyzed for each time point (n = 3 per time point). Body lengths were measured from the captured images.

### RNA extraction and cDNA synthesis

Total RNA was extracted from 5-dpf *ybx1*+/+ and *ybx1*−/− larvae using TRI-Reagent (Molecular Research Center, Cincinnati, OH) following the manufacturer’s protocol and as previously described ^17,18^. To eliminate genomic DNA contamination, RNA samples (10 *μ*g RNA in a 100 *μ*L reaction volume) were treated with DNase I (2U; New England Biolabs, Ipswich, MA) for 10 min at 37°C. The DNase-treated RNA was then purified by phenol-chloroform extraction and ethanol precipitation. Reverse transcription (RT) was performed using 0.5 µg of oligo(dT) primer, 1X M-MLV RT buffer, 0.5 mM each dNTP, 0.1 mM dithiothreitol (DTT), and 100 U of M-MLV reverse transcriptase (Invitrogen, Thermo Fisher Scientific, CA) in a 10 µL reaction volume at 37°C for 2 h. For quantitative comparisons, 3 µg of total RNA from each sample was used for reverse transcription (RT). RNA integrity was assessed using the Agilent Bioanalyzer 2100 (Agilent, Stockport, UK).

### RNA-sequencing and data analysis

RNA-seq libraries were constructed using the NEBNext Ultra Directional RNA Library Prep Kit (New England Biolabs) according to the manufacturer’s instructions. Libraries were sequenced on the Illumina HiSeq 2500 Sequencing System (Illumina, San Diego, CA) to generate 100-bp paired-end reads. RNA-seq data analysis was performed at the Genomics and Bioinformatics Core (Faculty of Health Sciences, University of Macau). Raw sequencing data have been deposited in the NCBI Gene Expression Omnibus (GEO) database under accession number GSE134334. Differential gene expression, Gene Ontology (GO) enrichment, and Kyoto Encyclopedia of Genes and Genomes (KEGG) pathway enrichment analyses were conducted as previously described ^19^.

### Real-time quantitative PCR (RT-qPCR)

To validate RNA-seq data, expression levels of selected genes (*nox1, duox, alpi2, fabp2, fabp6, mmp9* and *mmp13a,*) were quantified in 5-dpf *ybx1*+/+ and *ybx1*−/− larvae using RT-qPCR, with normalization to *ef1a* expression. Gene expression was also assessed in adult *ybx1*+/+ and *ybx1*−/− intestines. Additionally, *mmp9* and *mmp13a* mRNA levels were measured in *ybx1*−/− larvae at 5, 10, 15, and 20 dpf following exposure to ascorbic acid (vitamin C, VC) and hydrogen peroxide (H_2_O_2_). Furthermore, twelve inflammation-related genes reported ^20^ were analyzed by RT-qPCR. Primers for each gene are listed in Supplementary Table S1. RT-qPCR was performed using the CFX96 Real-Time PCR Detection System (Bio-Rad).

### Immunoblotting

WT embryos (*ybx1*+/+) were collected and maintained post-spawning. Larvae at various developmental stages (1-10 dpf, or between 5.00 and 6.25 dpf at finer intervals: 5.00, 5.17, 5.25, 5.33, 5.50, 5.67, 6.00, and 6.17 dpf) and those treated with 50 µM MG132 (5-6 dpf) were lysed in SDS sample buffer [62.5 mM Tris-HCl pH 6.8, 1% (w/v) SDS, 10% glycerol, 5 β-mercaptoethanol] (20 larvae/400 mL). Lysates were heated (95°C, 10 min), centrifuged (15 min, 4°C), and protein concentration determined with the 2-D Quant kit (Sigma-Aldrich, St. Louis, MO). Protein samples (100 µg) were separated by SDS-PAGE (12.5% polyacrylamide gels) and transferred to PVDF membranes. Membranes were blocked with 5% non-fat milk in TBST at room temperature (RT, 1 h) and incubated overnight at 4°C with primary antibodies against Ybx1 (1:2000 dilution) ^12^ and β-actin (1:2000 dilution) (Rabbit mAb, #4970; Cell Signaling, Danvers, MA) in 5% non-fat milk in TBST. After washing three times in TBST, membranes were incubated at RT for 1 h with HRP-conjugated secondary antibody (1:2000 dilution in TBST). After washing three times in TBST, protein bands were visualized using the SuperSignal West Femto Maximum Sensitivity Substrate (Thermo Scientific, Waltham, MA) and imaged on the ChemiDoc MP imaging system (Bio-Rad, Hercules, CA).

### Cold stress experiment

WT embryos were maintained at 28 ± 1°C. At 3 dpf, larvae were transferred to a glass bottle and subjected to cold stress at 16°C in a refrigerated water bath (MRC Lab, Cambridge, UK). Larvae were collected at 4, 5, and 6 dpf for protein extraction, as described above. This experiment was performed in duplicate. Ybx1 protein levels were quantified by Western blotting (as described above) in larvae maintained at 28°C (control) and those exposed to 16°C, at each time point (4, 5, and 6 dpf).

### Diethylaminobenzaldehyde treatment

N,N-diethylaminobenzaldehyde (DEAB), an inhibitor of aldehyde dehydrogenases (ALDHs) ^21^, was used to investigate the relationship between intestinal differentiation and Ybx1 expression. Previous studies in zebrafish have shown that treatment with DEAB or knockdown of retinal dehydrogenase (RDH), a homolog of human ALDHs, impairs intestinal differentiation ^22,23^. At 3 dpf (52 hpf), WT larvae (*ybx1*+/+) were divided into two groups: a control group and a DEAB-treated group. The DEAB-treated group was exposed to 100 µM DEAB, while the control group received no DEAB. Larvae from both groups were collected at 4 dpf (76 hpf), 5 dpf (100 hpf), and 6 dpf (126 hpf). For each time point and group, half of the larvae were fixed for histological examination, and the other half were used for protein preparation and Western blot analysis (as described above). The DEAB treatment experiment was performed in duplicate.

### Immunohistochemical (IHC) and immunofluorescent (IF) staining

The *ybx1*+/+ larvae at 5 and 6 dpf were fixed in Bouin’s solution overnight at 4°C. Fixed larvae were paraffin-embedded, sectioned (5 µm), and mounted on Superfrost Plus slides. Sections were deparaffinized, rehydrated, and subjected to antigen retrieval (10 mM sodium citrate buffer, 95°C, 10 min). Endogenous peroxidase activity was quenched with 3% H_2_O_2_ (10 min). After washing three times in PBS (5 min each), sections were blocked with normal horse serum (1 h, RT) and incubated overnight at 4°C with Ybx1 primary antibody (1:2000 dilution in blocking solution). Sections were washed (3x PBS) and incubated with secondary antibody (1 h, RT). Signal was amplified using the VECTASTAIN ABC-HRP Kit (Vector Laboratories, Burlingame, CA) (30 min, RT) and visualized with 3,3’-diaminobenzidine (DAB) (Vector Laboratories) (5 min). The reaction was stopped with running tap water, and slides dehydrated and mounted with Permount.

For immunofluorescent staining, the *ybx1*+/+ larvae at 5 dpf were fixed in 4% paraformaldehyde (PFA). Fixed larvae were paraffin-embedded, sectioned, deparaffinized, rehydrated, and subjected to antigen retrieval as described above. After serum blocking (as described above), sections were incubated overnight at 4°C with anti-Ybx1 primary antibody (1:2000 dilution in blocking solution). Sections were then incubated with Alexa Fluor 488- or Alexa Fluor 568-conjugated secondary antibody (Life Technologies, Carlsbad, CA). Nuclei were counterstained with DAPI. Slides were mounted and imaged using a Nikon ECLIPSE Ni-U microscope (Nikon).

### Histological examination

For histological analysis, *ybx1*+/+ and *ybx1*−/− larvae (5 dpf) and fish at various developmental stages (10, 20, 30, 45, 60, 90, 120, 150, 180, and 240 dpf) were fixed in Bouin’s solution overnight at RT. Fixed samples were dehydrated, paraffin-embedded, and serially sectioned (7 µm). Sections were stained with hematoxylin and eosin (H&E) and imaged using the ECLIPSE Ni-U microscope (Nikon).

### Whole mount in situ hybridization (WISH)

WISH was performed to examine the spatial expression of *nox1, duox, alpi2, fabp2, fabp6, mmp9* and *mmp13a*. The mRNA probes were transcribed in vitro and stored at −80°C. The WISH protocol was adapted from a previous study ^24^ with modifications. Briefly, 5-dpf *ybx1*+/+ and *ybx1*−/− larvae were fixed in 4% PFA overnight at 4°C, washed in PBST, and permeabilized with proteinase K (10 µg/mL, 15 min, RT). Samples were refixed (4% PFA, 1 h, RT), washed, and prehybridized (hybridization mix, 2 h, 65°C). Hybridization with digoxigenin-labeled RNA probes (100 ng in hybridization mix) was performed at 65°C for 40 h. Post-hybridization washes included graded series of hybridization mix/2x SSC, 2x SSC, 0.2x SSC + 0.1% Tween-20, and 0.1x SSC + 0.1% Tween-20 at 65°C, followed by washes in graded series of 0.2x SSC/PBST and PBST at RT. Samples were blocked in PBST with 2% sheep serum and 2 mg/mL BSA (1 h, RT) and incubated overnight at 4°C with anti-digoxigenin antibody conjugated to alkaline phosphatase (1:5000 dilution) (Merck, Darmstadt, Germany). After washing in PBST, color development was performed using NBT/BCIP in coloration buffer. The reaction was stopped with PBST (pH 5.5). Samples were mounted in glycerol and imaged using the SMZ18 microscope (Nikon).

### Enzyme-linked immunosorbent assay (ELISA)

Protein levels of alkaline phosphatase, intestinal (ALPI) were quantified in lysates from 5-dpf *ybx1*+/+ and *ybx1*−/− larvae and adult *ybx1*+/+ and *ybx1*−/− intestines. Samples were homogenized in ELISA lysis buffer (100 mM Tris, 150 mM NaCl, 1 mM EGTA, 1 mM EDTA, 1% Triton X-100, 0.5% Sodium deoxycholate), and protein concentrations were determined with the 2-D Quant kit (Sigma-Aldrich). ALPI levels were measured using the ALPI ELISA kit (OKEH00324, Aviva System Biology, San Diego, CA) according to the manufacturer’s instructions. Briefly, standards and diluted samples (100 µL, triplicate) were added to the ALPI Microplate and incubated (2 h, 37°C). After washing, Biotinylated ALPI Detector Antibody (100 µL, 1x) was added and incubated (1 h, 37°C). Following washing (3x), Avidin-HRP Conjugate (100 µL, 1x) was added and incubated (1 h, 37°C). Plates were washed (5x), and TMB Substrate (90 µL) was added. After incubation (15-30 min, 37°C, dark), Stop Solution (50 µL) was added. Absorbance was measured at 450 nm within 5 min using a microplate reader.

### Total antioxidant capacity (TAC) assay

Intestines from *ybx1*+/+ and *ybx1*−/− zebrafish were dissected, with samples pooled from three individuals per group (three groups in total for each genotype). Samples were homogenized in cold PBS containing EDTA-free protease inhibitor cocktail (Roche, Berlin, Germany) and centrifuged (10,000 g, 10 min, 4°C). Supernatants were aliquoted and stored at −80°C. Protein concentration was determined using the 2-D Quant kit (Sigma-Aldrich). Total antioxidant capacity was measured using the Total Antioxidant Capacity (TAC) Assay Kit (MBS168358, MyBioSource, San Diego, CA). Diluted uric acid standards and samples (20 µL, triplicate) were added to a 96-well plate, followed by 1x Reaction Buffer (180 µL). Initial absorbance was read at 490 nm. The reaction was initiated by adding 1x Copper Ion Reagent (50 µL) and incubating for 5 min with shaking. The reaction was terminated with 1x Stop Solution (50 µL), and the final absorbance was read at 490 nm. TAC was calculated based on the absorbance values.

### In vitro reactive oxygen species (ROS) assay

The ROS assay was performed using the In Vitro ROS/RNS Assay Kit (MBS168257, MyBioSource). All reagents were prepared and mixed before use. Samples, including intestine samples from *ybx1*+/+ and *ybx1*−/− zebrafish, and H_2_O_2_ standards were assayed in triplicate. Samples or standards (50 µL each) were added to a 96-well fluorescence plate, followed by 50 µL of catalyst. After mixing and a 5-min incubation at RT, 100 µL of DCFH solution was added. The plate was covered, incubated at RT for 15-45 min, and read on a fluorescence plate reader (480 nm excitation/530 nm emission). ROS levels were calculated using the fluorescence values and the H_2_O_2_ standard curve.

### Treatments with ascorbic acid and hydrogen peroxide

Since reduced antioxidant capacity and increased oxidative stress could be a factor for the high mortality in *ybx1*−/− zebrafish, we tested this by treating larvae and adults with ascorbic acid (VC) and hydrogen peroxide (H₂O₂), respectively. Briefly, the offspring of the *ybx1*+/− crosses were treated with 50 mg/L VC from 6 to 20 dpf, with daily water changes and fresh VC added each time. The treatment was designed with reference to previous studies in other fish species ^25,26^. H₂O₂ was used to enhance ROS stress according to a previous study ^27^. Based on a 24-h survival assay with *ybx1*+/+ larvae exposed to different concentrations of H₂O₂ (0.1, 0.5, 1.0, 1.5, and 2 mM), the concentrations of 0.1 and 0.5 mM H₂O₂ were selected for 15-day continuous exposure (6-20 dpf), with daily water changes and fresh H₂O₂ added. Control groups included larvae with normal feeding and larvae fed with only paramecia. For genotyping, 40 larvae were collected at 5, 10, 15, and 20 dpf (three parallel groups). Identified *ybx1*−/−larvae (5, 10, 15, and 20 dpf) from untreated, VC-treated, and H₂O₂-treated groups were used for RNA preparation. For adult treatment, the adult *ybx1*+/+ and *ybx1*−/− fish were treated with 50 mg/L VC or 0.5 mM H₂O₂ for 14 days, with daily water changes and fresh solutions. After treatment, fish were fixed for histological examination (three groups per treatment, 3-4 fish per group).

### Treatments with MMP9 and MMP13 inhibitors

To investigate the causal relationship between *mmp9/mmp13a* upregulation and postnatal larval lethality in *ybx1*−/− zebrafish, larvae and adults were treated with MMP inhibitors. The larvae from *ybx1*+/− crosses were cultured in Petri dishes, fed with paramecia, and treated from 6 to 20 dpf with MMP9 Inhibitor II (200 µM), MMP13 Inhibitor (20 µM), or MMP9/13 Inhibitor I (50 µM) (Merck Millipore, Burlington, MA), which were added to the water. Larvae were collected at 5 (pre-treatment), 10, 15, and 20 dpf for genotyping (40 individuals per time point, three parallel groups). For adults, the *ybx1*+/+ and *ybx1*−/− fish were treated with the same concentrations of MMP9 Inhibitor II, MMP13 Inhibitor, or MMP9/13 Inhibitor I for 14 days. After treatment, fish were fixed for histological examination (three groups per treatment, 3-5 fish per group).

### Observation and quantification of neutrophils in ybx1^−/−^ larvae intestine

To determine if intestinal inflammation was present in *ybx1*−/− larvae, *ybx1*−/−zebrafish were crossed with an EGFP-labeled neutrophil/macrophage transgenic line *Tg(coro1a:eGFP)hkz04t* obtained from China Zebrafish Resource Center (CZRC) ^28^ to generate *Tg(coro1a:eGFP)hkz04t;ybx1*−/−. The transgenic fish larvae were imaged using the SMZ18 microscope (Nikon). Additionally, intestinal tissue sections from both genotypes were imaged using the ECLIPSE Ni-U microscope (Nikon) to visualize EGFP fluorescence.

### Statistics

Target gene mRNA levels were normalized to the internal control *ef1a*. In some cases, data were further normalized as a percentage or fold change relative to a control or reference group. Data are presented as mean ± SEM. Statistical comparisons were performed using unpaired Student’s t-tests (GraphPad Prism 6, San Diego, CA).

## Results

### Effects of Ybx1 deficiency on perinatal and postnatal lethality

We observed maternal-effect lethality in the offspring of female *ybx1*−/− (maternal mutant M*ybx1*−/−). Most embryos from the M*ybx1*−/− females showed abnormal development with smaller sizes as indicated by reduced weight and diameter at 24 hours post-fertilization (hpf) (Fig. 1A-C). Most of these embryos died before hatching. Some hatched but died within a week before 7 dpf, with over 60% exhibiting severe abnormalities such as cardiac edema (Fig. 1D-F), resulting in 100% perinatal or postnatal lethality (Fig. 1G). In comparison, larvae from control females (*ybx1*+/+) developed well.

**Figure 1.**
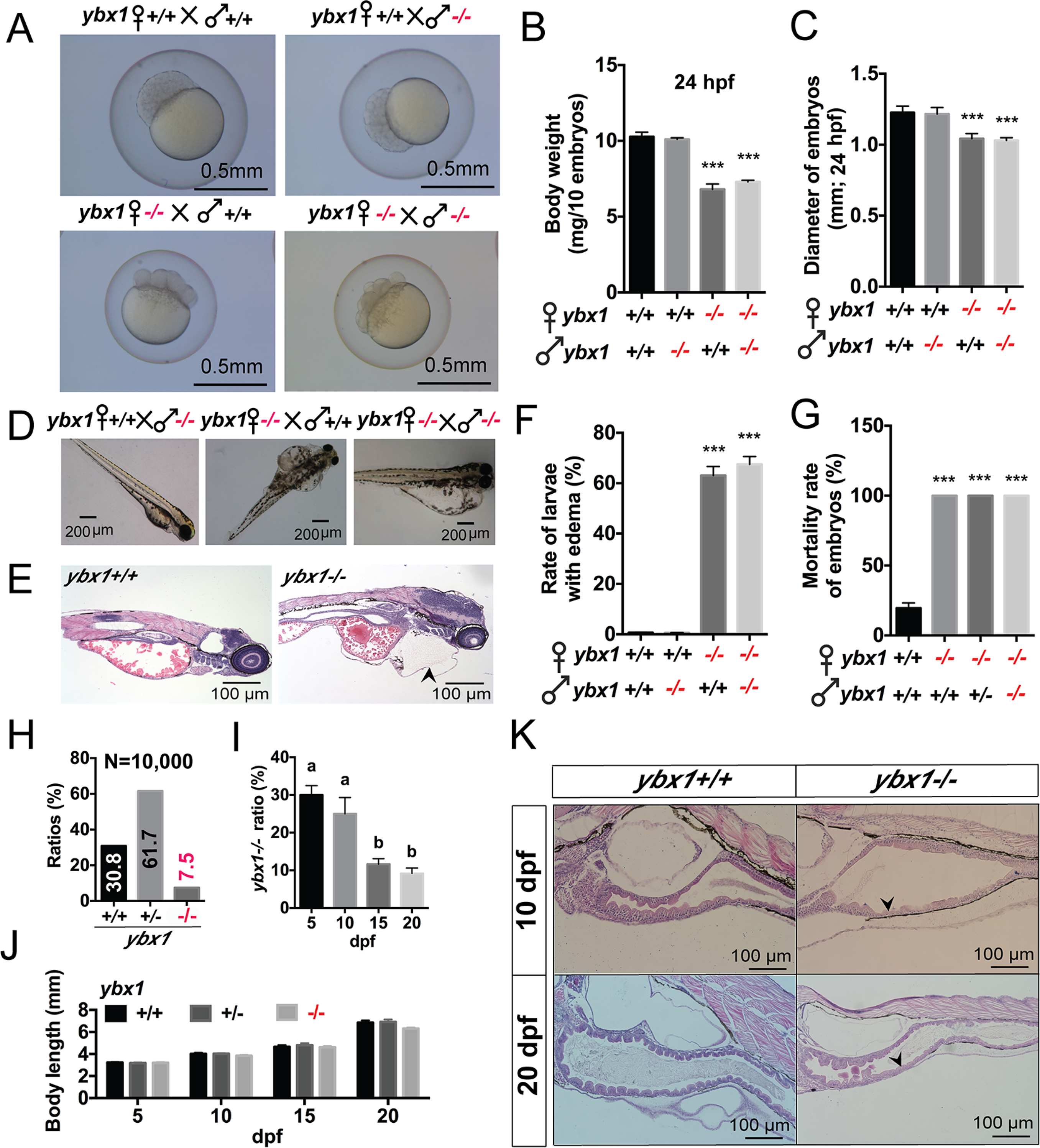
Maternal and zygotic deficiency of Ybx1 leads to distinct developmental defects. (**A–C**) Maternal Ybx1 is essential for early embryonic development. **(A)** Representative images of early embryos derived from the indicated crosses. Offspring from *ybx1* mutant females (*ybx1−/−* ♀) exhibited developmental delays regardless of the paternal genotype. **(B** and **C)** Quantification of embryonic body weight (10 embryos per measurement) (**B**) and diameter (**C**) at 24 hpf. Offspring from mutant females are significantly smaller and lighter. **(D–G)** Maternal-effect lethality in *ybx1* mutants. **(D)** Representative morphology of larvae at 72 hpf showing severe cardiac/abdominal edema in offspring of *ybx1−/−* females. **(E)** H&E staining of sagittal sections of 72-hpf larvae. The arrowhead indicates severe edema in the *ybx1−/−* larva. **(F)** Quantification of edema rates and **(G)** mortality rates. Offspring from *ybx1−/−* females show 100% mortality. Data are presented as mean ± SEM. ***p < 0.001. **(H–K)** Zygotic *ybx1* loss leads to partial postnatal lethality and intestinal defects. **(H)** Genotype distribution of progeny from heterozygous intercrosses (N = 10,000), showing a sub-Mendelian ratio (7.5%) of *ybx1−/−* survivors. **(I)** Survival analysis showing a significant drop in the ratio of *ybx1−/−* mutants between 10 and 15 dpf. Different letters (a, b) indicate statistical significance (p < 0.05). **(J)** Body length measurements at 5, 10, 15, and 20 dpf showing no significant growth retardation in surviving mutants. **(K)** H&E staining of intestinal sections at 10 and 20 dpf. Arrowheads indicate intestinal folds (villi). Note the thinning of the intestinal wall and shortening of villi in *ybx1−/−* mutants at 20 dpf.

In addition to maternal lethality, we also observed postnatal lethality in *ybx1*-deficient zygotic mutant (Z*ybx1*−/−). In the offspring derived from heterozygous parents (*ybx1*+/−), we found a significantly reduced proportion of *ybx1*−/− adults. Genotyping of 10,000 fish in total from heterozygous crosses identified 3082 *ybx1*+/+ (30.8%), 6165 *ybx1*+/− (61.7%), and 753 *ybx1*−/− (7.5%; ∼30% of homozygous mutant *ybx1*−/−) (Fig. 1H).

To determine the timing of Z*ybx1*−/− lethality, we performed genotyping at 5, 10, 15, and 20 dpf. The *ybx1*−/− ratio was within the expected Mendelian range at 5 and 10 dpf but decreased sharply at 15 and 20 dpf (Fig. 1I). This indicates that the critical period for post-hatching or postnatal lethality of *ybx1*−/− mutant larvae was between 10 and 15 dpf. Although the average body length of *ybx1*−/− larvae was slightly smaller at various time points (Fig. 1J), histological examination revealed a pronounced difference in intestinal development. Specifically, the overall intestinal surface in the mutant (*ybx1*−/−) appeared flatter than that of the age-matched control fish (*ybx1*+/+); in particular, the mucosal folds or villi of mutant intestines were shorter, fewer in number, and simpler in structure at both 10 and 20 dpf (Fig. 1K), suggesting impaired intestinal development.

### Dynamic regulation of Ybx1 expression during larval development

To understand the mechanism by which *ybx1* deficiency leads to postnatal mortality, we examined the spatiotemporal expression patterns of Ybx1 protein in WT larvae by Western blotting and IHC/IF staining. The larval samples were collected daily from 1 to 10 dpf for protein extraction. Interestingly, Ybx1 protein was expressed within a specific time window, appearing at 3 dpf and peaking at 5 dpf. However, this was followed by a sharp decrease to undetectable level at 6 dpf (Fig. 2A). This rapid and marked decline was attributed to protein degradation via the ubiquitin-proteasome system, as treatment with the proteasome inhibitor MG132 prevented Ybx1 degradation at 6 dpf (Fig. 2B). To further characterize this degradation, we determined Ybx1 levels at more frequent intervals between 5 and 6 dpf (Fig. S1). IHC and IF staining of 5 and 6-dpf larvae localized Ybx1 predominantly to the intestinal enterocytes (Fig. 2C and D). The observed degradation of Ybx1 at 6 dpf and its predominant intestinal localization led us to hypothesize a role for Ybx1 in intestinal development, as proposed above based on intestinal morphology.

**Figure 2.**
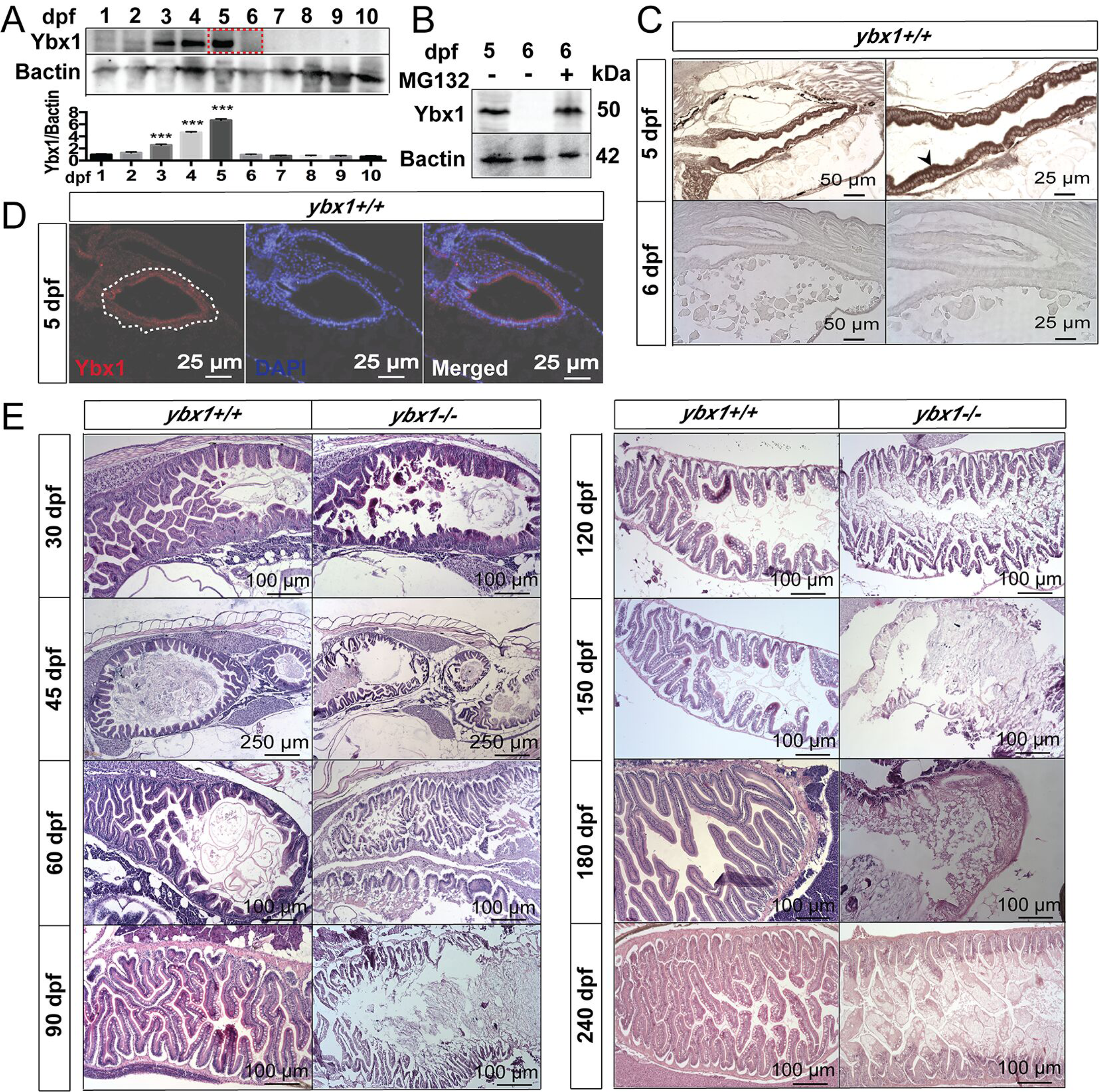
Spatiotemporal expression profiles of Ybx1 in embryos and postnatal larvae and its association with intestinal development. **(A)** Western blot analysis of Ybx1 protein levels in WT embryos and larvae from 1 to 10 dpf. Ybx1 expression peaked at 5 dpf and decreased sharply at 6 dpf (highlighted by the red dashed box). Quantification of the Ybx1/Bactin (β-actin) ratio is shown below. Data are presented as mean ± SEM. ***p < 0.001. **(B)** Western blot analysis of Ybx1 protein levels at 5 and 6 dpf, with and without treatment with the proteasome inhibitor MG132. Treatment with MG132 stabilized Ybx1 levels at 6 dpf, indicating degradation via the ubiquitin-proteasome pathway. **(C)** IHC staining of Ybx1 in *ybx1+/+* larvae. At 5 dpf, Ybx1 was strongly expressed in the intestinal epithelium (arrowhead), whereas signal was largely absent by 6 dpf. **(D)** IF staining for Ybx1 in sections of *ybx1+/+* larva. Ybx1 (red) was localized to the intestinal epithelium. Nuclei are stained with DAPI (blue). The white dashed line outlines the intestine. **(E)** H&E staining of intestinal sections from *ybx1+/+* and *ybx1−/−* fish from juvenile to adult stages (30, 45, 60, 90, 120, 150, 180, and 240 dpf). While WT fish maintained normal and healthy intestinal folds, *ybx1−/−* mutants displayed progressive intestinal defects, including blunted folds and severe tissue atrophy.

To provide further evidence for the association between the temporal expression pattern of Ybx1 protein and specific stages of embryonic development, we performed two experiments. First, we reduced the incubation temperature from 28°C to 16°C at 3 dpf to decelerate the developmental process. Larval samples were collected from both control (28°C) and treatment (16°C) groups at 4, 5, and 6 dpf, and Ybx1 protein levels were assessed using immunoblotting. In the 16°C group, robust Ybx1 expression persisted through 6 dpf; in contrast, Ybx1 was undetectable in 6 dpf larvae maintained at 28°C (Fig. S2). Second, we treated the larvae with 100 µM diethylaminobenzaldehyde (DEAB), an inhibitor of aldehyde dehydrogenases (ALDH) known to inhibit intestinal differentiation ^23^, and examined Ybx1 expression at 4, 5, and 6 dpf. Both control and DEAB-treated larvae were collected at these time points for histological analysis and immunoblotting. DEAB treatment markedly inhibited intestinal differentiation and development, resulting in flattened surfaces with shortened villi at 6 dpf (Fig. S3A), in contrast to the normal villus morphology observed in untreated age-matched larvae. Immunoblotting revealed persistent Ybx1 expression in DEAB-treated larvae at 6 dpf (126 hpf), whereas Ybx1 was undetectable in the control group at this time point (Fig. S3B). The sustained Ybx1 expression in larvae was closely associated with the impaired intestinal development following DEAB treatment, suggesting a critical role for Ybx1 degradation at 6 dpf in intestinal differentiation.

### Long-term effects of Ybx1 deficiency on intestinal integrity

To assess the long-term effects of Ybx1 deficiency, we performed histological analysis on *ybx1*+/+ and *ybx1*−/− zebrafish at various stages from juvenile to adult (30, 45, 60, 90, 120, 150, 180, and 240 dpf). Intestinal villus defects were already apparent in *ybx1*−/− zebrafish at 30 dpf, and these defects progressively worsened as the fish grew. In early stages from 30 to 60 dpf, the control fish (*ybx1*+/+) showed normal intestinal architecture with well-shaped villi, well-organized epithelial lining, clear lumen and consistent mucosal folds. In contrast, the intestines of mutant (*ybx1*−/−) displayed a serious disruption of villus structure with irregular shapes and reduced height. The lumen appeared dilated with compromised mucosal integrity. Also, epithelial degeneration started to appear with obvious signs of epithelial atrophy. In intermediate stages from 90 to 150 pdf, the intestinal structure remained consistent in the control fish, with tightly packed and elongated villi; whereas the mutant intestines showed severe disorganization with loss of villus integrity, collapse of mucosal structures, and signs of tissue degeneration. In late stages from 180 to 240 dpf, the control intestines have fully developed with robust mucosal lining. However, the mutant intestines had lost normal architecture with extensive tissue fibrosis and severe epithelial degeneration (Fig. 2E). These phenotypes of intestinal abnormalities in *ybx1*−/− fish resemble features of certain human gastrointestinal disorders, especially the inflammatory bowel disease (IBD), which is characteristic of chronic inflammation, mucosal damage, epithelial destruction, and fibrosis ^29^.

### Implication of oxidative stress in intestinal dysfunction of ybx1 mutants

To investigate the molecular mechanisms underlying postnatal lethality in *ybx1*−/−zebrafish, we performed an RNA-seq analysis on 5-dpf larvae when Ybx1 protein reached its peak level. Transcriptomic comparison of *ybx1*+/+ and *ybx1*−/− larvae revealed numerous differentially expressed genes (DEGs), as visualized in the heatmap (Fig. 3A) and volcano plot (Fig. 3B). GO enrichment analysis revealed a significant upregulation of pathways associated with inflammation and oxidative stress. Several GO terms related to inflammation were upregulated in *ybx1*−/− larvae, including inflammatory response, response to bacterium, response to cytokines and macrophage chemotaxis; also significantly upregulated was the term reactive oxygen species biosynthetic process (Fig. 3C and D). KEGG enrichment analysis revealed significant upregulation of genes associated with apoptosis, lysosome, and phagosome pathways (Fig. 3E).

**Figure 3.**
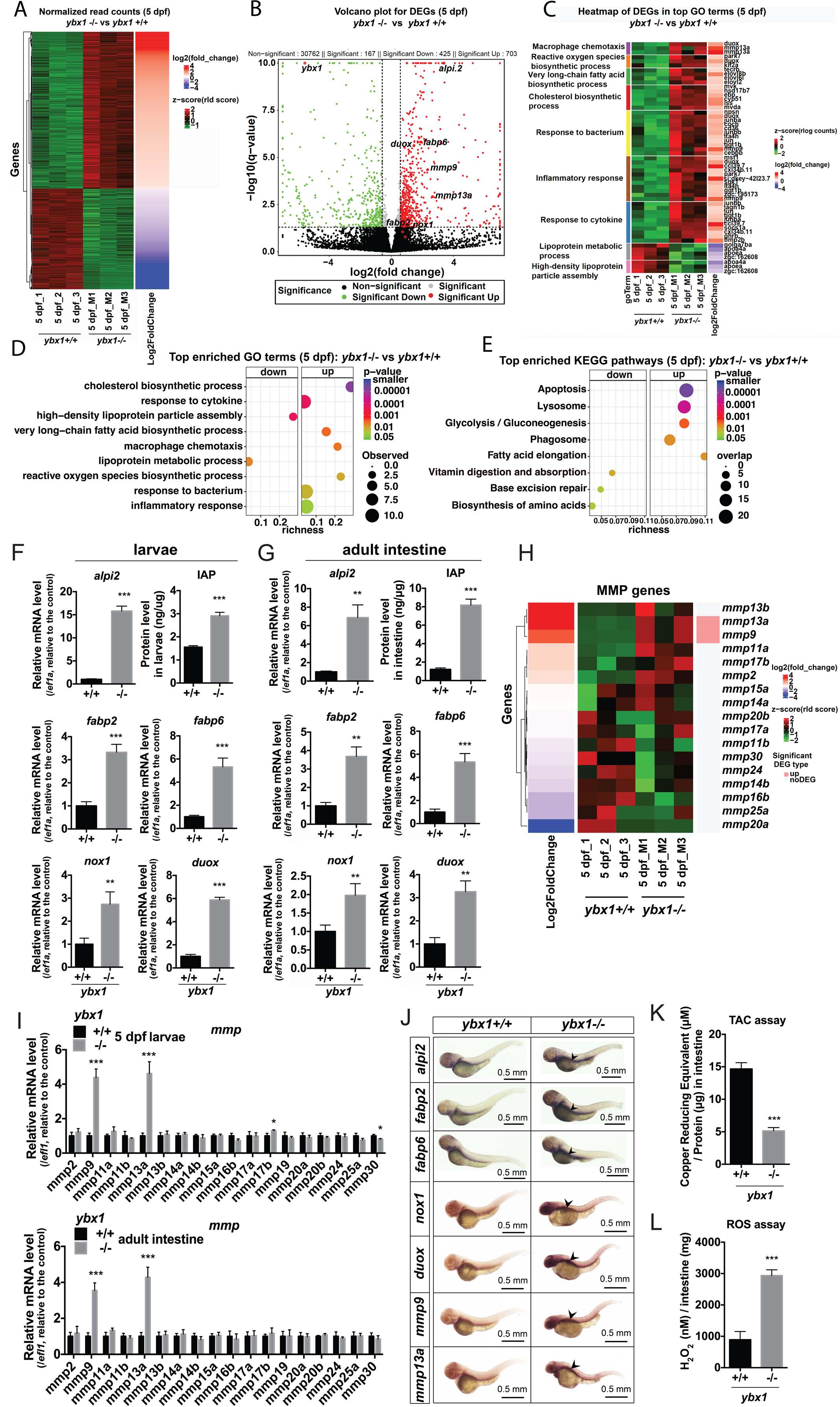
Transcriptomic analysis revealing upregulation of inflammatory, oxidative stress, and differentiation markers in *ybx1* mutants. **(A)** Heatmap of differentially expressed genes (DEGs) comparing *ybx1+/+* and *ybx1−/−* larvae at 5 dpf (n=3 biological replicates per group). **(B)** Volcano plot illustrating DEGs. Red and green dots indicate significantly upregulated and downregulated genes, respectively. Key genes involved in inflammation (*mmp9, mmp13a*), ROS production (*nox1, duox*), and intestinal differentiation (*alpi.2, fabp2, fabp6*) are labeled. **(C)** Heatmap of DEGs clustered by Gene Ontology (GO) terms. Upregulated clusters include inflammatory response, response to bacterium, response to cytokine, and ROS biosynthetic processes, while downregulated clusters involve lipid metabolism. **(D)** Dot plot showing the top enriched GO terms. Upregulated pathways (right) are dominated by immune response and ROS production, while downregulated pathways (left) are related to lipid and cholesterol metabolism. **(E)** KEGG pathway enrichment analysis showing upregulation of apoptosis, lysosome and phagosome pathways, and downregulation of metabolic pathways like biosynthesis of amino acids. **(F** and **G)** Validation of RNA-seq data by RT-qPCR (mRNA) and biochemical assay (IAP protein) in 5-dpf larvae **(F)** and adult intestines **(G)**. Expression levels of *alpi2, fabp2, fabp6, nox1,* and *duox* are significantly elevated in mutants. IAP protein levels are also significantly increased. **(H)** Heatmap specifically displaying the expression profiles of matrix metalloproteinase (*mmp*) family genes. **(I)** RT-qPCR analysis of *mmp* genes in 5-dpf larvae (top) and adult intestines (bottom). The *mmp9* and *mmp13a* genes were consistently and significantly upregulated in *ybx1−/−* mutants compared to *ybx1+/+* controls. **(J)** Whole-mount in situ hybridization of 5-dpf larvae. Stronger signals for *alpi2, fabp2, fabp6, nox1, duox, mmp9,* and *mmp13a* were observed in the intestines of *ybx1−/−* larvae (black arrowheads) compared to wild-type siblings. **(K)** Total Antioxidant Capacity (TAC) assay showing significantly reduced antioxidant capacity in the intestines of *ybx1−/−* fish. **(L)** ROS assay showing significantly elevated oxidative stress (H_2_O_2_) levels in *ybx1−/−* intestines. Data are presented as mean ± SEM. *P<0.05, **p < 0.01, ***p < 0.001.

To validate the RNA-seq data, we performed RT-qPCR on selected DEGs in both whole larvae and dissected intestines from *ybx1*+/+ and *ybx1*−/− zebrafish. The selected genes included intestinal alkaline phosphatase 2 (*alpi2*), fatty acid-binding protein 2 (*fabp2*), fatty acid-binding protein 6 (*fabp6*), NADPH oxidase (*nox1*), dual oxidase (*duox*), and 18 members of the MMP family. These genes were chosen because they are either specifically expressed in the intestine (*alpi2*, *fabp2,* and *fabp6*) or associated with ROS production (*nox1* and *duox*) and tissue remodeling (MMP family). RT-qPCR analysis confirmed the upregulation of *alpi2* in both *ybx1*−/− larvae and adult intestines. This was further confirmed by ELISA, which demonstrated a significant increase in intestinal alkaline phosphatase (IAP) activity in *ybx1*−/− samples. The expression patterns of *fabp2, fabp6*, *nox1* and *duox* were also consistent with the RNA-seq data (Fig. 3F, G). Additionally, among the 18 MMP family members examined, *mmp9* and *mmp13a* showed a marked upregulation in both *ybx1*−/− larvae and adult intestines (Fig. 3H, I). Overall, the RT-qPCR results validated the RNA-seq findings.

To determine the spatial distribution of the aforementioned DEGs, we performed whole-mount in situ hybridization (WISH) on *ybx1*+/+ and *ybx1*−/− larvae at 5 dpf. We focused on *alpi2, fabp2, fabp6, nox1, duox, mmp9* and *mmp13a*. The result showed predominant gut localization of *nox1, duox, mmp9* and *mmp13a* mRNA, with increased expression in *ybx1*−/− larvae. The expression of *alpi2, fabp2*, and *fabp6* served as positive controls, as their gut-specific expression has been reported in humans ^30–32^. In zebrafish larvae, WISH confirmed the predominant gut localization of *alpi2, fabp2*, and *fabp6* transcripts, also with significantly increased expression in *ybx1*−/− larvae (Fig. 3J).

Consistent with the observation of increased reactive oxygen species biosynthetic process in the mutant, we found that the total antioxidant capacity (TAC) was significantly reduced in the intestines of *ybx1*−/− zebrafish compared to *ybx1*+/+ (Fig. 3K). Furthermore, we observed a marked elevation of reactive oxygen species (ROS) levels in the intestines of *ybx1*−/− zebrafish (Fig. 3L). These findings suggest that increased ROS levels and reduced antioxidant capacity in the intestine may contribute to the observed intestinal defects and postnatal lethality in *ybx1*−/− zebrafish.

### Effects of exogenous antioxidant and prooxidant on postnatal survival and adult intestinal integrity

The postnatal lethality in the *ybx1*−/− mutant occurred between 10 and 15 dpf, coinciding with the diet change from paramecia to artemia in our aquarium system. To determine if this diet change was associated with the death of mutant fish, we tested two feeding schemes: one with the standard diet transition and another with continuous paramecia feeding (Fig. 4A). The results showed no significant difference in lethality between the two groups (Fig. 4B).

**Figure 4.**
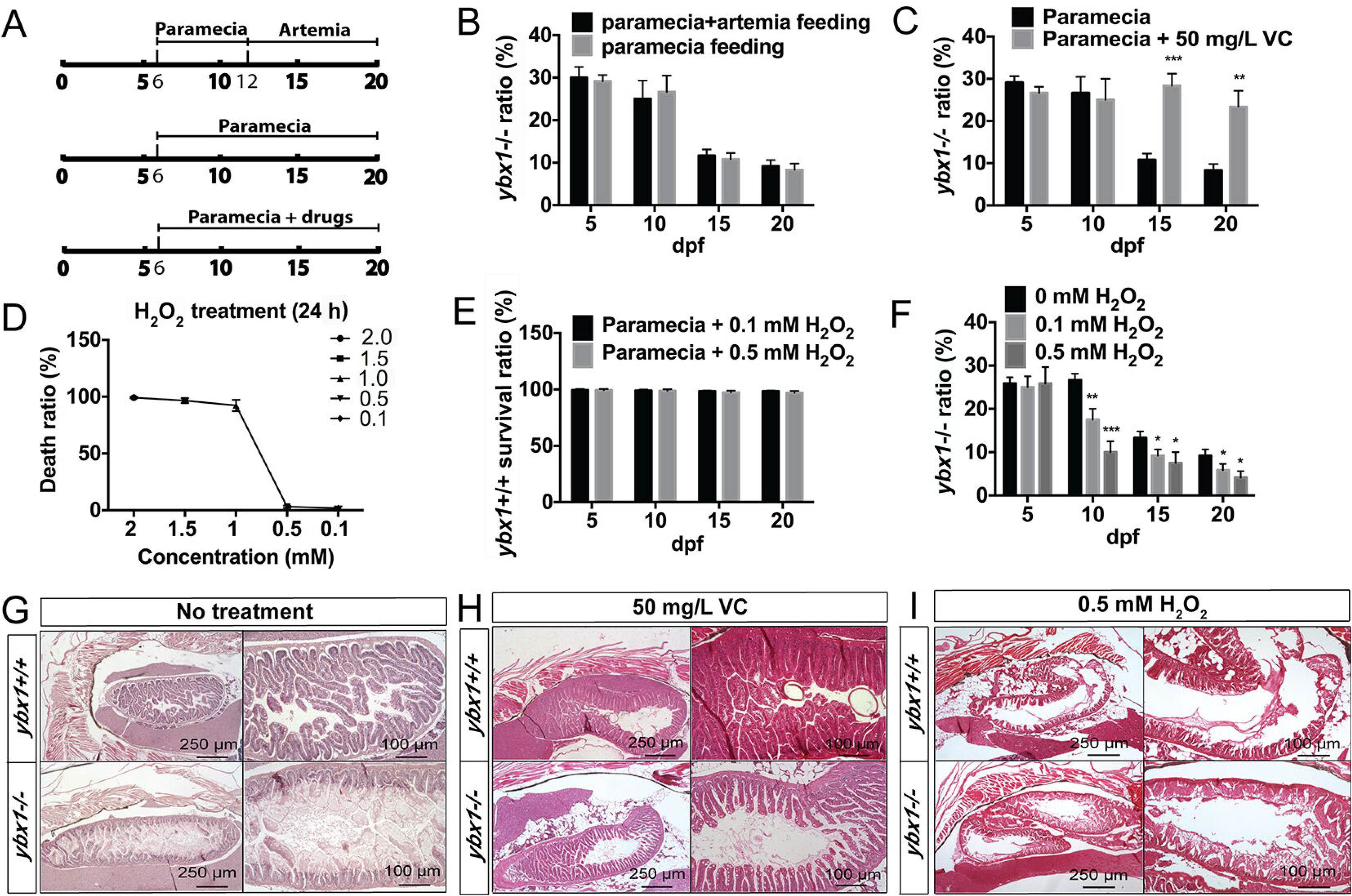
Involvement of oxidative stress in intestinal defects and lethality in *ybx1*−/− mutants. (**A**) Schematic diagram of the experimental timelines for feeding and drug administration. Larvae were fed paramecia starting at 6 dpf, with some groups switching to artemia at 12 dpf. Drug treatments (VC or H_2_O_2_) were conducted with paramecia feeding. (**B**) Survival analysis (ratio of *ybx1−/−* mutants) comparing larvae fed only paramecia versus those switched to artemia. Changing the diet to artemia did not affect the lethality of *ybx1−/−* mutants. (**C**) Survival analysis of *ybx1−/−* mutants treated with 50 mg/L VC. VC treatment significantly rescued the death phenotype of the mutants at 15 and 20 dpf compared to untreated controls. (**D**) Dose-response curve for H_2_O_2_ toxicity in WT larvae treated for 24 h. Concentrations of 0.1 and 0.5 mM resulted in 0% mortality and were selected for sublethal exposure experiments. (**E**) Survival rate of WT larvae treated with 0.1 or 0.5 mM H_2_O_2_ from 5 to 20 dpf. These concentrations had no effect on WT survival. (**F**) Survival analysis of *ybx1*−/− mutants under oxidative stress. Treatment with 0.1 or 0.5 mM H_2_O_2_ significantly accelerated the decline in the mutant ratio as early as 10 dpf compared to the 0 mM control, indicating hypersensitivity to oxidative stress. (**G–I**) Histological analysis of adult intestines from *ybx1*+/+ and *ybx1*−/− fish under different conditions. (**G)** Control fish without drug treatment. The *ybx1*−/− intestines showed disrupted mucosal folds and thinner walls compared to *ybx1*+/+ control. (**H**) Treatment with VC (50 mg/L) rescued the intestinal morphology in *ybx1*−/− mutants, restoring fold structure. (**I**) Treatment with H_2_O_2_ (0.5 mM) exacerbated the intestinal damage in *ybx1*−/− mutants, leading to severe tissue disintegration compared to *ybx1*+/+ control. Data are presented as mean ± SEM. *p < 0.05, **p < 0.01, ***p < 0.001.

To investigate the role of oxidative stress in *ybx1*−/− postnatal lethality, we first examined the effect of ascorbic acid (vitamin C, VC), a well-known antioxidant, on the postnatal lethality in the mutant. Treatment with VC (50 mg/L) successfully rescued the lethal phenotype of the mutant fish (*ybx1*−/−) at 15 and 20 dpf, with a normal Mendelian ratio of approximately 25% (Fig. 4C). To further assess the impact of oxidative stress, we exposed fish larvae to hydrogen peroxide (H_2_O_2_), a potent prooxidant, at concentrations of 0.1 and 0.5 mM, which were doses that did not affect fish survival (Fig. 4D, E). As expected, treatment with H_2_O_2_ induced significant lethality at 10 dpf in a dose-dependent manner, and it further reduced the mutant survival rates at 15 and 20 dpf (Fig. 4F), indicating that increased ROS levels exacerbate postnatal lethality.

We then extended our investigation to adult stage to examine how oxidative stress affected intestinal structure. We first treated *ybx1*+/+ and *ybx1*−/− fish (∼120 dpf) with 50 mg/L VC for two weeks, changing the water and replenishing VC daily. Control group received no VC treatment. Following treatment, fish were processed for histological examination. In treatment control group, the intestine of the wild-type fish (*ybx1*+/+) appeared healthy and intact, displaying a highly organized structure with villi densely packed and lined by a uniform layer of epithelial cells. In contrast, the intestine of the *ybx1−/−* mutant fish showed severe structural disorganization. The villi were significantly shorter, blunted, and irregularly shaped. The intricate folded structure is largely lost, resulting in a deteriorating and likely dysfunctional intestinal lining. There appeared to be significant amount of cellular debris within the intestinal lumen, suggesting widespread cell death and sloughing of the epithelial lining (Fig. 4G). This observation of tissue degradation is consistent with phenotypes caused by severe oxidative stress ^33^. In VC treatment group, the intestine of *ybx1*+/+ fish appeared the same as the untreated wild-type fish. The villi were long, well-organized, and the tissue looked healthy. This suggests that under normal conditions, exposure to VC did not significantly alter the intestinal structure at the concentration used (50 mg/L). The effect of VC on the *ybx1−/−* mutant fish was remarkable. The intestinal structure showed a significant recovery compared to the untreated mutant in the control group. The villi were more elongated and better defined, resembling the wild-type structure. While not a complete restoration, the overall organization was vastly improved, and there is a visible reduction in luminal debris (Fig. 4H). This rescue effect by an antioxidant strongly supports the hypothesis that oxidative stress, at least in part, contributed to the mutant phenotype. In contrast to the treatment with antioxidant, the exposure to prooxidant (H_2_O_2_) showed opposite effects. The H_2_O_2_-treated WT fish (*ybx1+/+*) showed signs of intestine damage, with tissue structure being less regular and villi appearing shorter and somewhat disorganized compared to the untreated fish. This indicates that exposure to this level of H_2_O_2_ was toxic and could induce oxidative stress even in healthy individuals. In *ybx1−/−* mutant fish, H_2_O_2_ treatment exacerbated the phenotype of the mutant. The intestinal structure was almost completely disintegrated. The villi were virtually absent, replaced by a flattened and severely damaged epithelial layer. There was extensive tissue degradation with cellular debris and signs of inflammation (Fig. 4I). This severe worsening of the phenotype by a prooxidant provides compelling evidence that *ybx1* mutation made the intestinal tissue highly vulnerable to oxidative stress.

### Involvement of MMP9 and MMP13a in maintenance of intestinal integrity

Our transcriptome analysis and RT-qPCR assays revealed a remarkable up-regulation of *mmp9* and *mmp13a* expression among MMP family members in *ybx1*−/−mutant compared to the control fish (Fig. 3). To investigate the involvement of *mmp9* and *mmp13a* upregulation in *ybx1*−/− postnatal lethality and intestinal development, we treated *ybx1*+/+ and *ybx1*−/− larvae with three different MMP inhibitors: MMP9 inhibitor II, MMP13 inhibitor, and MMP9/13 inhibitor I. Treatments with 200 µM MMP9 inhibitor II, 20 µM MMP13 inhibitor and 50 µM MMP9/13 inhibitor I all markedly increased the survival rates of the mutant fish at 15 and 20 dpf (Fig. 5A-C). These results indicate that inhibition of MMP9 and MMP13 activity could rescue *ybx1*−/− postnatal lethality during the critical period of 10–20 dpf.

**Figure 5.**
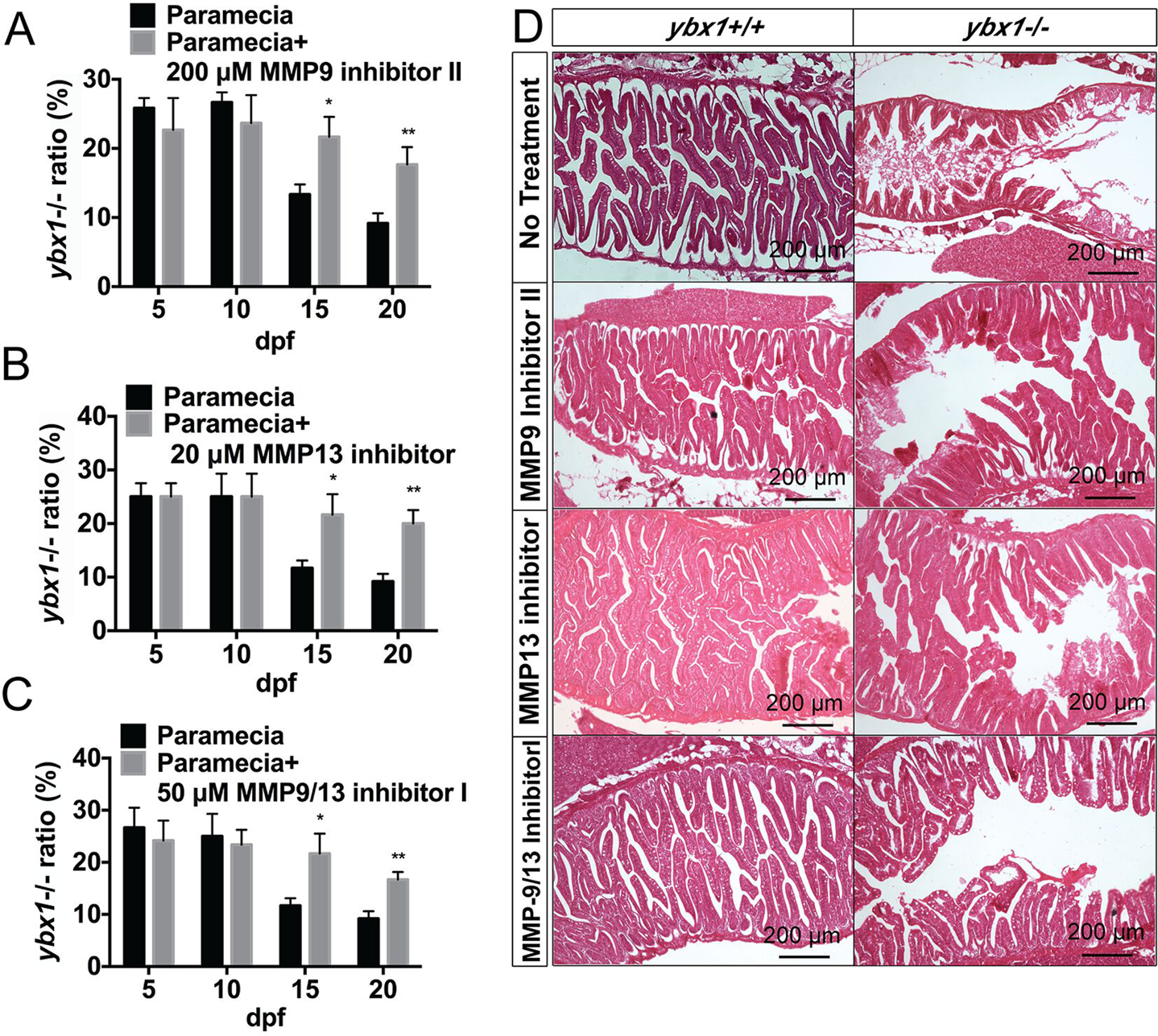
Effects of MMP9 and MMP13 inhibitors on lethality and intestinal defects in *ybx1*−/− mutants. (**A–C**) Survival analysis of *ybx1*−/− larvae treated with specific MMP inhibitors. (**A**) Effect of 200 µM MMP9 inhibitor II. (**B**) Effect of 20 µM MMP13 inhibitor. (**C**) Effect of 50 µM MMP9/13 inhibitor I. All three inhibitors could significantly increase the survival rate of *ybx1*−/− larvae. (**D**) Histological examination of intestines from *ybx1*+/+ and *ybx1*−/− larvae at 20 dpf under different treatment conditions. The top row shows that untreated *ybx1*−/− mutants exhibited severe intestinal atrophy and loss of mucosal fold structure compared to WT siblings. The rows below demonstrate that treatment with MMP9 inhibitor II, MMP13 inhibitor, or MMP9/13 inhibitor I effectively restored intestinal morphology and structural integrity in the *ybx1*−/− mutants. Data are presented as mean ± SEM. *p < 0.05, **p < 0.01.

To determine whether the rescue of mutant lethality by MMP9 and MMP13 inhibitors was related to intestinal development, we performed histological analysis at the end of the treatment. As described above, the mutant intestine displayed a severe pathology without inhibitor treatment. The villi were dramatically shortened, blunted, and fused, with a disorganized epithelial lining. The lumen contained significant amount of cellular debris, suggesting that the intestinal epithelial cells were dying and sloughing off, leading to a catastrophic failure of tissue integrity. Treatments of the mutant fish with the MMP9 and MMP13 inhibitors alone both resulted in a noticeable improvement of the intestinal phenotype. Compared to the untreated mutant, the villi were more elongated and better defined, and the overall structure was better organized. While the villi were still shorter and less complex than in the *ybx1*+/+ fish, the severe tissue degradation was clearly reduced. This partial rescue indicates that MMP9 and MMP13 activities were important contributors to the intestinal damage. Simultaneous inhibition of both MMP9 and MMP13 activities with MMP9/13 inhibitor showed the most dramatic and effective rescue. The intestinal structure in the mutant fish treated with the dual MMP-9/13 inhibitor was substantially improved. The villi were much longer, more abundant, and better organized in a complex folded pattern that closely resembles the wild-type structure. While perhaps not a complete restoration, the improvement appeared greater than that seen with either the MMP9 or MMP13 inhibitor alone (Fig. 5D).

### ROS regulation of MMP9 and MMP13a expression

To investigate whether the elevated expression of *mmp9* and *mmp13a* in *ybx1−/−*mutants is regulated by oxidative stress, we performed RT-qPCR to measure their expression levels in fish of different genotypes (*ybx1+/+*, *ybx1+/−*, and *ybx1−/−*) and their responses to either H₂O₂ (0.5 mM) or VC (50 mg/L) treatment at multiple time points post-fertilization.

First, we examined the effect of prooxidant H₂O₂ on MMP gene expression. In *ybx1−/−* mutant fish, exposure to H₂O₂ resulted in a significant and time-dependent upregulation of both *mmp9* and *mmp13a* mRNA compared to untreated mutants fed with paramecia (Fig. 6A, B). A statistically significant increase in *mmp9* expression was first detected at 10 dpf (p<0.01), with the effect becoming more pronounced at 15 and 20 dpf, ultimately resulting in an approximate 4-fold elevation compared to control levels (p<0.001) (Fig. 6A). A similar pattern was observed for *mmp13a*, with expression increasing significantly from 10 dpf onwards and peaking at 15 and 20 dpf with an approximately 3-fold increase (p<0.001) (Fig. 6B). To determine if this response was specific to the mutant background, we compared the effect of H₂O₂ across all three genotypes at 5 and 20 dpf. At 5 dpf, H₂O₂ exposure did not cause significant changes in *mmp9* or *mmp13a* expression in any genotype. However, at 20 dpf, a significant genotype-dependent effect was observed. H₂O₂ dramatically increased *mmp9* or *mmp13a* expression by approximately 4.5-fold in *ybx1−/−* mutants, but not in the control fish (*ybx1+/+* and *ybx1+/−*) (Fig. 6C, D). This demonstrates that *ybx1−/−*mutants are hypersensitive to oxidative stress, which leads to a profound overexpression of these matrix metalloproteinases.

**Figure 6.**
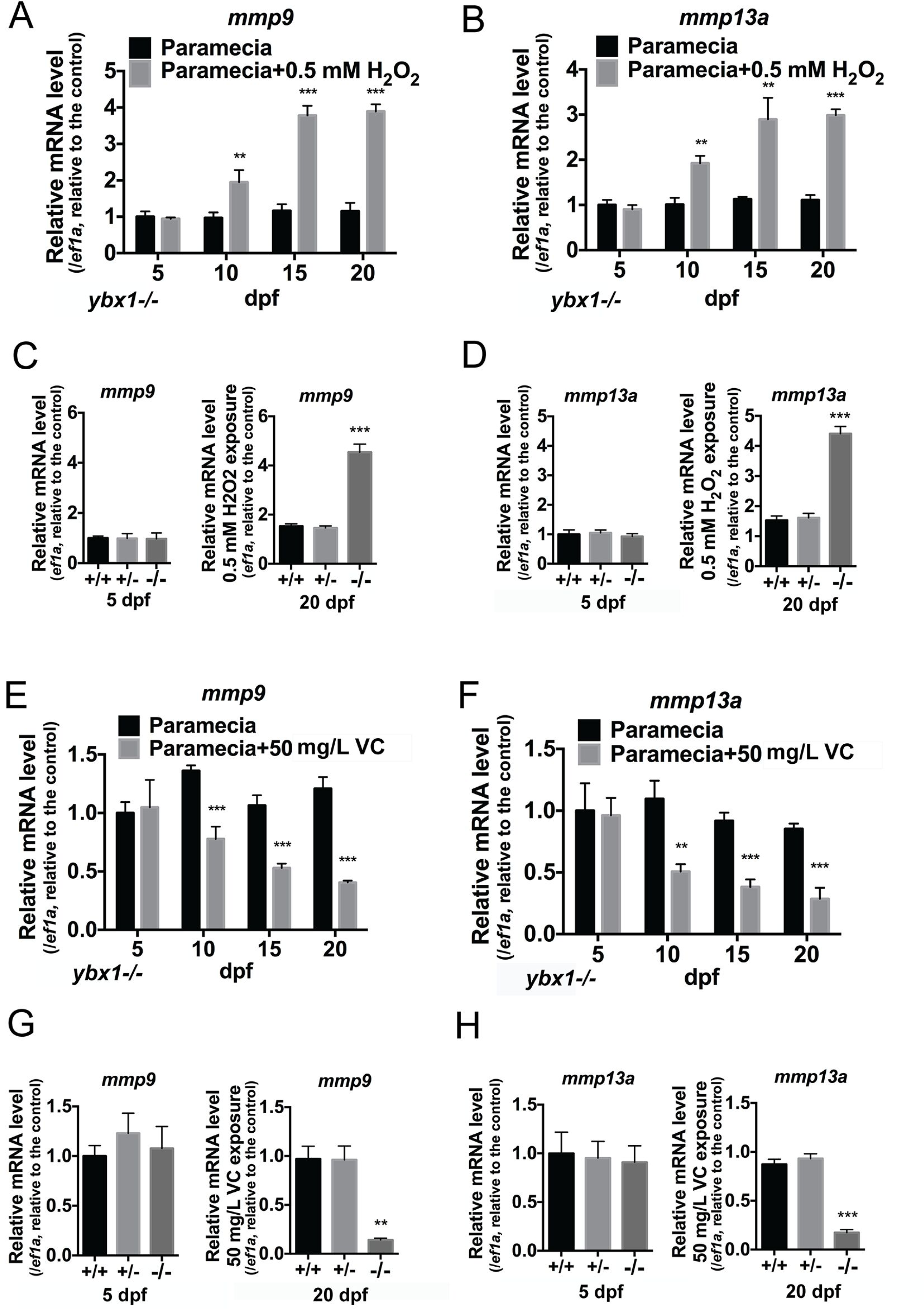
Responses of *mmp9* and *mmp13a* expression to prooxidant and antioxidant in *ybx1*−/− mutants. **(A** and **B)** Relative mRNA levels of *mmp9* (**A**) and *mmp13a* (**B**) in *ybx1−/−* larvae fed with paramecia (control) or paramecia supplemented with 0.5 mM H₂O₂ from 5 to 20 dpf. Exposure to H₂O₂ significantly increased the expression of both MMPs in the mutants from 10 dpf onwards. **(C** and **D)** Comparison of *mmp9* (**C**) and *mmp13a* (**D**) expression levels among *ybx1+/+*, *ybx1+/−*, and *ybx1−/−*siblings at 5 dpf (left panels) and at 20 dpf following 0.5 mM H₂O₂ exposure (right panels). At 20 dpf, H₂O₂ treatment induced a sharp increase in MMP expression specifically in the *ybx1−/−* mutants compared to WT and heterozygous siblings (*ybx1+/+*, *ybx1+/−*). **(E** and **F)** Relative mRNA levels of *mmp9* (**E**) and *mmp13a* (**F**) in *ybx1−/−* larvae treated with 50 mg/L VC. Treatment with VC significantly suppressed the mRNA levels of both genes in the mutants from 10 to 20 dpf compared to controls. **(G** and **H)** Comparison of *mmp9* (**G**) and *mmp13a* (**H**) expression levels among genotypes at 5 dpf (left panels) and 20 dpf (right panels) with 50 mg/L VC exposure. VC treatment resulted in significantly lower MMP expression in *ybx1−/−* mutants at 20 dpf. Gene expression levels were normalized to *ef1a*. Data are presented as mean ± SEM. **p < 0.01, ***p < 0.001.

We then investigated whether antioxidant could reverse the effect. We treated *ybx1−/−* mutants with VC and compared them to the untreated paramecia-fed group (Fig. 6E, F). The results showed that VC treatment led to a significant, time-dependent suppression of both *mmp9* and *mmp13a* expression. For both genes, a significant reduction was observed from 10 dpf onwards, with expression levels decreasing to less than 50% of the untreated controls by 20 dpf. This indicates that alleviating oxidative stress can effectively reduce the expression of these genes in the mutant background. We also compared the effect of VC exposure across all genotypes at 5 and 20 dpf. At 5 dpf, VC had no significant effect on *mmp9* or *mmp13a* expression in any genotype. At 20 dpf, VC treatment caused a dramatic and significant reduction in both *mmp9* and *mmp13a* mRNA levels in the mutant (*ybx1−/−*) compared to the control fish (*ybx1+/+* and *ybx1+/−*) (Fig. 6G, H). These observations indicate that *mmp9* and *mmp13a* expression in *ybx1*−/− larvae is highly sensitive to ROS levels, suggesting that these genes are ROS-inducible in the absence of Ybx1.

### Evidence for intestinal inflammation in ybx1−/− mutant

The severe tissue degradation and ROS-induced upregulation of *mmp9* and *mmp13a* expression in *ybx1−/−* mutants, along with the RNA-seq data showing increased expression of genes related to immune and inflammatory responses, strongly suggested the presence of a pathological inflammatory response. To confirm this, we assessed the inflammatory status in the mutant by introducing *ybx1−/−* mutant in the transgenic reporter line *Tg(coro1a:eGFP)hkz04t*, which specifically labels neutrophils with GFP. Neutrophil infiltration is considered a classic hallmark of acute inflammation ^34,35^. Immunofluorescent analysis of intestinal sections at 5 dpf revealed an obvious difference between genotypes. In *ybx1+/+* larvae [*Tg(coro1a:eGFP)hkz04t;ybx1*+/+], the intestinal tissue was well-structured with only a few sparsely distributed GFP-positive neutrophils. In contrast, the intestines of *ybx1−/−* larvae [*Tg(coro1a:eGFP)hkz04t;ybx1*−/−] exhibited a significant infiltration of neutrophils. These immune cells were seen to accumulate specifically within the disorganized and damaged intestinal tissue, concentrating in the lamina propria and epithelium (Fig. 7A). Quantification of these cells confirmed the visual observations, demonstrating a statistically significant, approximately 3-fold increase in the number of neutrophils within the intestinal region of *ybx1−/−* larvae compared to their *ybx1+/+* siblings (p<0.01) (Fig. 7B).

**Figure 7.**
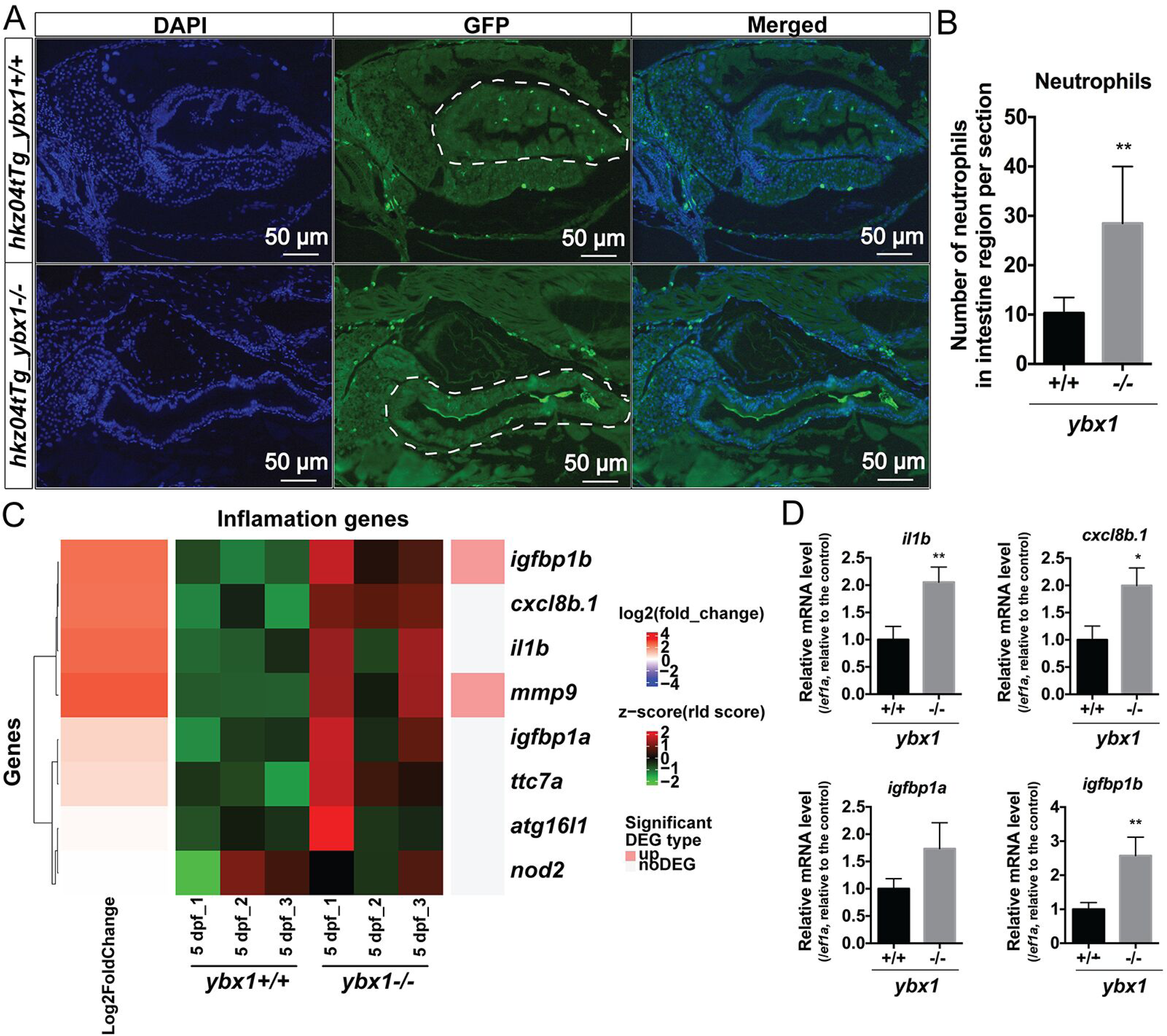
Intestinal inflammation and upregulation of inflammatory genes in *ybx1*−/− mutants at 5 dpf. **(A)** Representative immunofluorescent images of intestinal sections from *hkz04tTg_ybx1+/+* and *hkz04tTg_ybx1−/−* larvae. The *hkz04tTg* transgene marks neutrophils with GFP (green), and nuclei were stained with DAPI (blue). The white dashed lines outline the intestinal region. **(B)** Quantification of the number of neutrophils present in the intestine region per section. The mutants (*ybx1−/−*) showed a significantly higher number of infiltrating neutrophils compared to *ybx1+/+* siblings. **(C)** Heatmap illustrating differential expression of inflammation-related genes in *ybx1+/+* and *ybx1−/−* larvae at 5 dpf (n=3 biological replicates per genotype). The color scale represents the z-score (green = low expression, red = high expression). The sidebar indicates significant DEGs, with *igfbp1b* and *mmp9* highlighted as upregulated genes (pink) in the mutants. **(D)** Relative mRNA levels of inflammation-related genes (*il1b*, *cxcl8b.1*, *igfbp1a* and *igfbp1b*,) determined by RT-qPCR in *ybx1+/+* and *ybx1−/−*larvae. Expression levels were normalized to *ef1a*. Data are presented as mean ± SD. *p < 0.05, **p < 0.01.

To further confirm the intestinal inflammation in *ybx1−/−* larvae, we analyzed the expression of several pro-inflammatory genes (*il1b*, *cxcl8b.1, igfbp1a,* and *igfbp1b*). Consistent with the transcriptome data (Fig. 7C), RT-qPCR analysis at 5 dpf showed upregulation of mRNA levels of *il1b* (cytokine), *cxcl8b.1* (neutrophil chemoattractant), *igfbp1a* and *igfbp1b* (insulin-like growth factor binding protein-1a and b) in the mutant although the increase of *igfbp1a* was not statistically significant (Fig. 7D).

## Discussion

### Ybx1 is essential for embryonic and postnatal development

We demonstrated in this study that Ybx1 plays a critical role in zebrafish embryonic development, particularly in the development of intestine. Maternal *ybx1* deficiency (M*ybx1*−/−) resulted in severe perinatal lethality, characterized by phenotypes like cardiac edema and developmental retardation. Our observation of perinatal lethality from female mutants is consistent with previous zebrafish studies ^13,14^; additionally, we identified cardiac edema as a major phenotype of maternal *ybx1* deficiency.

In addition to perinatal lethality associated with maternal *ybx1* deficiency, we also observed postnatal lethality in zygotic *ybx1* mutants (Z*ybx1*−/−), with most deaths occurring between 10 and 20 dpf. This phenotype agrees well with previous studies in mice, where YB-1 deficiency caused embryonic lethality due to cellular stress sensitivity ^6^. Interestingly, the period of lethality was preceded by a three-day window (3 to 5 dpf) when the level of Ybx1 protein showed a dramatic surge, followed by a sharp decline to undetectable levels by 6 dpf. Further experiment showed that the disappearance of Ybx1 was due to protein degradation mediated by the ubiquitin-proteasome system. This is consistent with the reports in mammalian cells that YB-1 is a target protein for degradation by the ubiquitin-proteasome system ^36,37^. Interestingly, the zygotic mutant lethality exhibited incomplete genetic penetrance, as some mutant individuals were able to survive beyond the lethal window and reach adulthood. These results indicate that Ybx1 plays a significant role in postnatal development, and that the postnatal mortality observed in *ybx1−/−* mutants may be attributed not only to the loss of *ybx1* but also to additional internal and/or external factors.

### Ybx1 regulates intestinal differentiation and homeostasis

To understand the functional roles of Ybx1 in postnatal larvae, we performed IHC and IF staining to determine its localization of expression at 5 dpf when the protein level peaks. The results showed a strong presence of Ybx1 protein in the intestinal epithelium, which was not detected at 6 dpf. This change suggests a significant and specific role for Ybx1 in intestinal development during early larval stage. Pharmacological and environmental interventions, such as low-temperature stress and DEAB treatment, delayed Ybx1 degradation and hindered intestinal differentiation. These findings raise the intriguing possibility that Ybx1 acts as a gatekeeper or temporal brake on intestinal maturation, and its precise temporal expression and removal via the ubiquitin-proteasome system are required for terminal differentiation and maturation of enterocytes. In addition, *ybx1*−/− zebrafish displayed long-lasting defects in intestinal structure from juvenile to adult stages, indicating a crucial role for Ybx1 in maintaining intestinal homeostasis. To our knowledge, this study represents the first reporting involvement of YB-1 (Ybx1) in intestinal development and maintenance.

### Oxidative stress and intestinal inflammation in ybx1−/− zebrafish

RNA-seq analysis revealed that *ybx1*−/− larvae exhibited upregulation of genes involved in ROS biosynthesis and inflammatory response. This was consistent with reduced total antioxidant capacity and elevated ROS levels in mutant intestines. Specifically, *nox1* and *duox,* members of the NOX and DUOX families involved in ROS generation ^38,39^, were significantly upregulated in *ybx1*−/− larvae. Extensive studies have shown a close association between Nox1 and Doux levels and hydrogen peroxide (H_2_O_2_) production in vivo ^40–42^. The increased expression of *nox1* and *duox* in *ybx1*−/−zebrafish suggests that Ybx1 deficiency may lead to increased ROS production, which then drives intestinal inflammation and pathology. Indeed, histological analysis of *ybx1*−/− larvae showed that their intestines exhibited structural abnormalities associated with inflammation. Supporting this finding was an increased neutrophil presence within the intestine and higher expression levels of inflammatory markers such as *il1b*, *igfbp1b*, and *cxcl8b.1*. Together, these results point to an overall pro-inflammatory condition in the mutant intestine.

YB-1 is a multifunctional protein involved in various cellular functions, including responses to oxidative stress. It plays major roles in DNA and RNA repair, stress granule formation, cell survival, and regulation of oxidative damage. YB-1 stimulates DNA repair under oxidative stress by interacting with NEIL2, one of the four oxidized base-specific DNA glycosylases associated with the DNA base excision repair (BER) pathway responsible for removing damaged bases ^43^. It also binds oxidized mRNAs to prevent translation errors ^44,45^ and is recruited to stress granules (SGs) during oxidative stress to help sort and process damaged RNA ^46^. However, this mechanism does not seem to work in zebrafish cells, where YB-1-positive SGs can form in response to heat shock stress but not oxidative stress ^47^. Interestingly, oxidative stress also enhances YB-1 secretion, which can act as a paracrine/autocrine signal to induce cell cycle arrest and inhibit proliferation in neighboring cells, contributing to tissue-level stress responses ^46^. In mice, YB-1 mutant cells (−/−) exhibited reduced abilities to respond to oxidative stress and increased expression of cell cycle inhibitors such as p16 and p21, resulting in premature cell senescence ^6^. Recently, we also demonstrated that the loss of Ybx1 in zebrafish resulted in a surged expression of p21 in the mutant ovary, leading to arrested follicle development ^15^.

The expression data suggest a role for oxidative stress in the intestinal pathology and larval mortality of *ybx1−/−* zebrafish. This is further supported by results from two key experiments. First, antioxidant supplementation with ascorbic acid (VC) rescued *ybx1−/−* larvae from postnatal lethality and restored the intestinal villus structure in adult mutants. Second, increasing ROS levels through hydrogen peroxide (H_2_O_2_) exposure exacerbated both lethality and intestinal damage in a dose-dependent manner. These results implicate ROS levels as a critical determinant of *ybx1−/−* phenotypes and raise the possibility that antioxidant therapy could be effective in treating Ybx1-deficiency-related disorders.

### Roles of ROS-induced MMPs in intestinal pathology

Our findings that loss of *ybx1* leads to oxidative stress and subsequent upregulation of *mmp9* and *mmp13a* suggests a role for matrix metalloproteinases (MMPs) in zebrafish intestinal pathology observed in *ybx1*−/− mutant as MMPs are well known to play a critical role in tissue homeostasis and their disorders are often linked to various diseases ^48,49^. The role of Mmp9 and Mmp13a in intestinal damage and larval survival was further supported by our pharmacological experiment. Treatment of the *ybx1*−/−larvae with MMP9 and MMP13 inhibitors significantly improved the larval survival and intestinal morphology of the *ybx1*−/− mutant, indicating their critical role in the intestinal pathology associated with Ybx1 deficiency.

Under normal physiological conditions, the activities of MMPs are tightly controlled for physiological processes like tissue remodeling and wound healing ^50^. Specifically, MMP-13 (Collagenase-3) is a potent collagenase with strong ability to degrade fibrillar collagens, which are major components of extracellular matrix (ECM), while MMP-9 (Gelatinase B) specializes in degrading type IV collagen, a key component of basement membranes that anchor the epithelial layer ^51^. The delicate balance between ECM deposition and degradation is frequently disrupted in pathological conditions. Excessive MMP activity, as observed in our *ybx1−/−* model, leads to uncontrolled breakdown of the ECM, resulting in a catastrophic loss of tissue architecture and compromised epithelial barrier integrity, a hallmark of human inflammatory bowel diseases (IBD) ^52^.

A key aspect of MMP pathology is their regulation by cellular stress signals, particularly reactive oxygen species (ROS). It is well established that oxidative stress can potently induce the expression of MMP genes at the transcriptional level, often through the activation of redox-sensitive signaling pathways, especially MMP9 and MMP13 ^53–56^. Our data, showing that exposure to prooxidant H₂O₂ dramatically increased *mmp9* and *mmp13a* expression while the antioxidant vitamin C suppressed it, provides direct evidence for this regulatory axis in our model. Furthermore, MMPs are not merely downstream effectors of tissue damage; they are active participants and amplifiers of the inflammatory response. MMP-9, in particular, acts as a powerful pro-inflammatory mediator by processing and activating cytokines and chemokines, such as interleukin-8 (IL-8), a potent neutrophil chemoattractant that plays an important role in immune responses at mucosal sites ^57,58^. This creates a vicious feedback loop where initial tissue damage and ROS production trigger MMP expression, which in turn degrades tissue and promotes further immune cell infiltration, perpetuating the inflammatory cycle and exacerbating the pathology. The significant neutrophil recruitment we observed in the intestines of *ybx1−/−* larvae is likely a direct consequence of such ROS-MMP axis, providing a mechanistic link between the loss of Ybx1 and the onset of severe intestinal inflammation.

### Ybx1-deficient zebrafish as a model for ROS-induced inflammatory bowel disease

The phenotypes observed in *ybx1*-deficient zebrafish intestine, including the severe degradation of intestinal architecture, compromised epithelial barrier, and neutrophil-driven inflammation, present a mechanistically defined model for key aspects of human inflammatory bowel disease (IBD). The value of the *ybx1−/−* zebrafish model extends beyond phenotypical similarities. It recapitulates a pathogenic cascade potentially implicated in human IBD: a primary genetic defect leading to chronic oxidative stress, which in turn drives the pathological upregulation of matrix metalloproteinases and induces a destructive inflammatory response.

The inherent advantages of the zebrafish system, including its genetic tractability, optical transparency for high-resolution *in vivo* imaging of inflammatory processes, and suitability for rapid, high-throughput chemical screening, have been instrumental for disease modeling. Recent studies from other laboratories have also attempted to model IBD-like diseases in zebrafish, demonstrating its potential as a promising alternative to traditional murine models ^59,60^. The powerful reverse genetics in zebrafish has enabled creation of IBD-like models with symptoms resembling those in human patients. For example, morpholino-mediated knockdown of two IBD susceptibility genes nucleotide-binding oligomerization domain containing 1 and 2 (*nod1* and *nod2*), which are expressed in intestinal epithelial cells and neutrophils, reduced embryonic response to infection and decreased expression of *duox* in the intestinal epithelium ^61^. A recent study showed that mutation of class III PI3-kinaes (*pik3c3*) in zebrafish caused intestinal injury and inflammation, as evidenced by disrupted microvilli, loss of adherens junctions, and neutrophil infiltration ^20^. The *ybx1−/−* mutant described in this study exhibited all major characteristics of human IBD, making it another promising model for future research. It provides a much-needed *in vivo* platform to investigate how chronic oxidative stress initiates and perpetuates intestinal inflammation, to dissect the specific roles of MMPs in tissue destruction and immune signaling, and to screen for novel therapeutic compounds that can disrupt this vicious cycle. In addition, this model offers a unique opportunity to explore the complex interplay between genetic susceptibility and environmental stressors in the pathogenesis of intestinal disorders like IBD.

In summary, this study establishes Ybx1 as a critical regulator of embryonic and intestinal development, with its deficiency leading to ROS-driven intestinal inflammation and lethality. The rescue of *ybx1*−/− phenotypes by antioxidants and MMP inhibitors highlights potential therapeutic strategies for ROS-MMP axis-related disorders. Future studies should investigate the molecular mechanisms by which Ybx1 regulates ROS homeostasis and explore its role in other oxidative stress-related diseases. Additionally, the development of small-molecule therapies targeting ROS or MMP activity could offer new treatments for conditions similar to those involving Ybx1 dysfunction.

## Acknowledgements

We thank Ms. Phoenix Un Ian LEI for the maintenance and management of the zebrafish facility and the Genomics and Bioinformatics Core of the Faculty of Health Sciences for technical support. This project has been supported by CPG2024-00030-FHS, CPG2025-00037-FHS, MYRG2022-00219-FHS, MYRG-GRG2023-00144-FHS-UMDF and MYRG-GRG2024-00191-FHS from University of Macau, and FDCT-NSFC Joint Project 0086/2022/AFJ from The Macau Fund for Development of Science and Technology.

---

**Figure S1.**
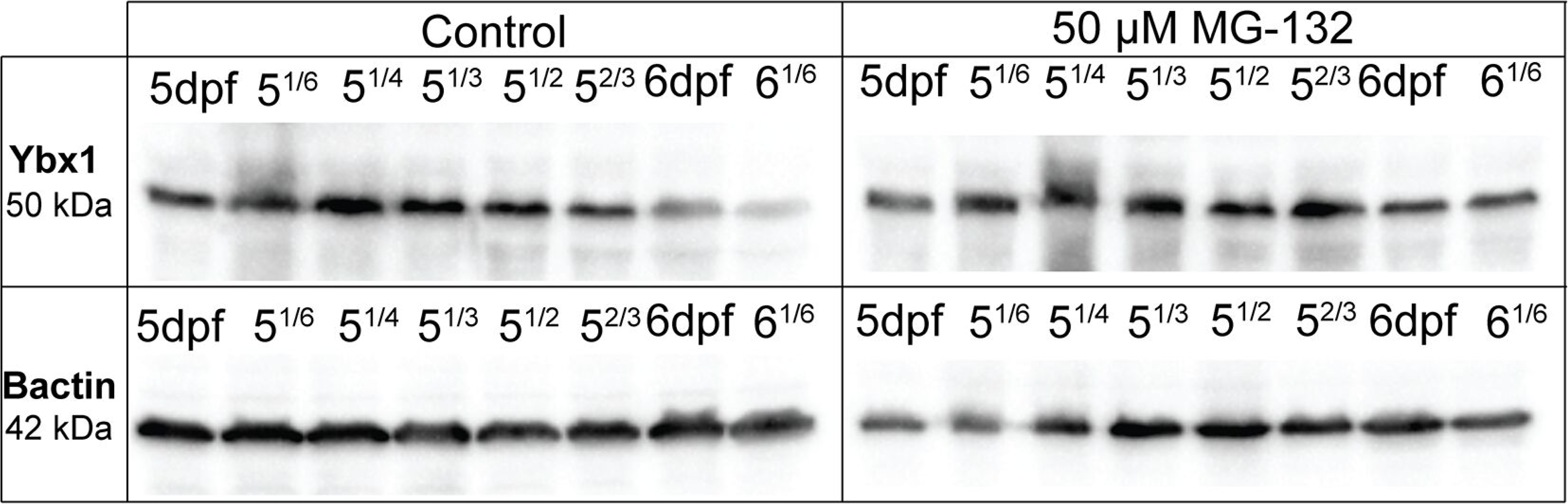
Stabilization of Ybx1 protein by proteasome inhibition. Western blot analysis of Ybx1 protein (50 kDa) abundance in zebrafish larvae treated with vehicle (Control) or the proteasome inhibitor MG-132 (50 µM). Samples were collected at various time points between 5 and 6 dpf (indicated as fractions of a day). The housekeeping gene (Bactin, 42 kDa) was used as the loading control. The stabilization and accumulation of Ybx1 in the MG-132 treated group compared to the control suggests that proteasomal degradation is a major mechanism for removing Ybx1.

**Figure S2.**
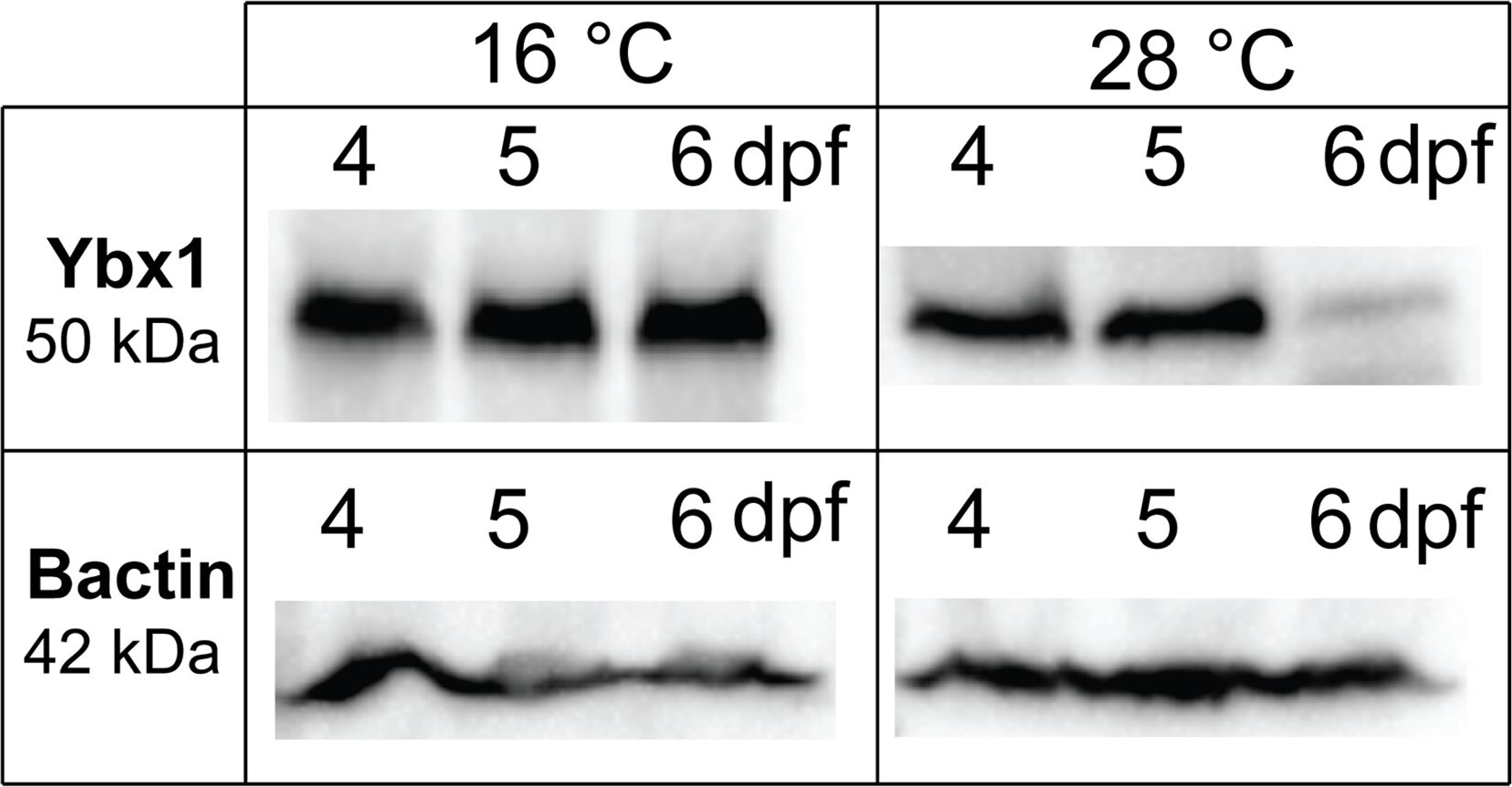
Temperature-dependent regulation of Ybx1 protein levels. Western blot analysis comparing Ybx1 (50 kDa) protein abundance in zebrafish larvae reared at a lower temperature (16°C) versus the standard temperature (28°C). Samples were collected at 4, 5, and 6 dpf. The housekeeping gene β-actin (Bactin, 42 kDa) was used as the loading control. The results show that while Ybx1 levels decreased significantly by 6 dpf at 28°C, the protein remained stable and abundant at 6 dpf when reared at 16°C.

**Figure S3.**
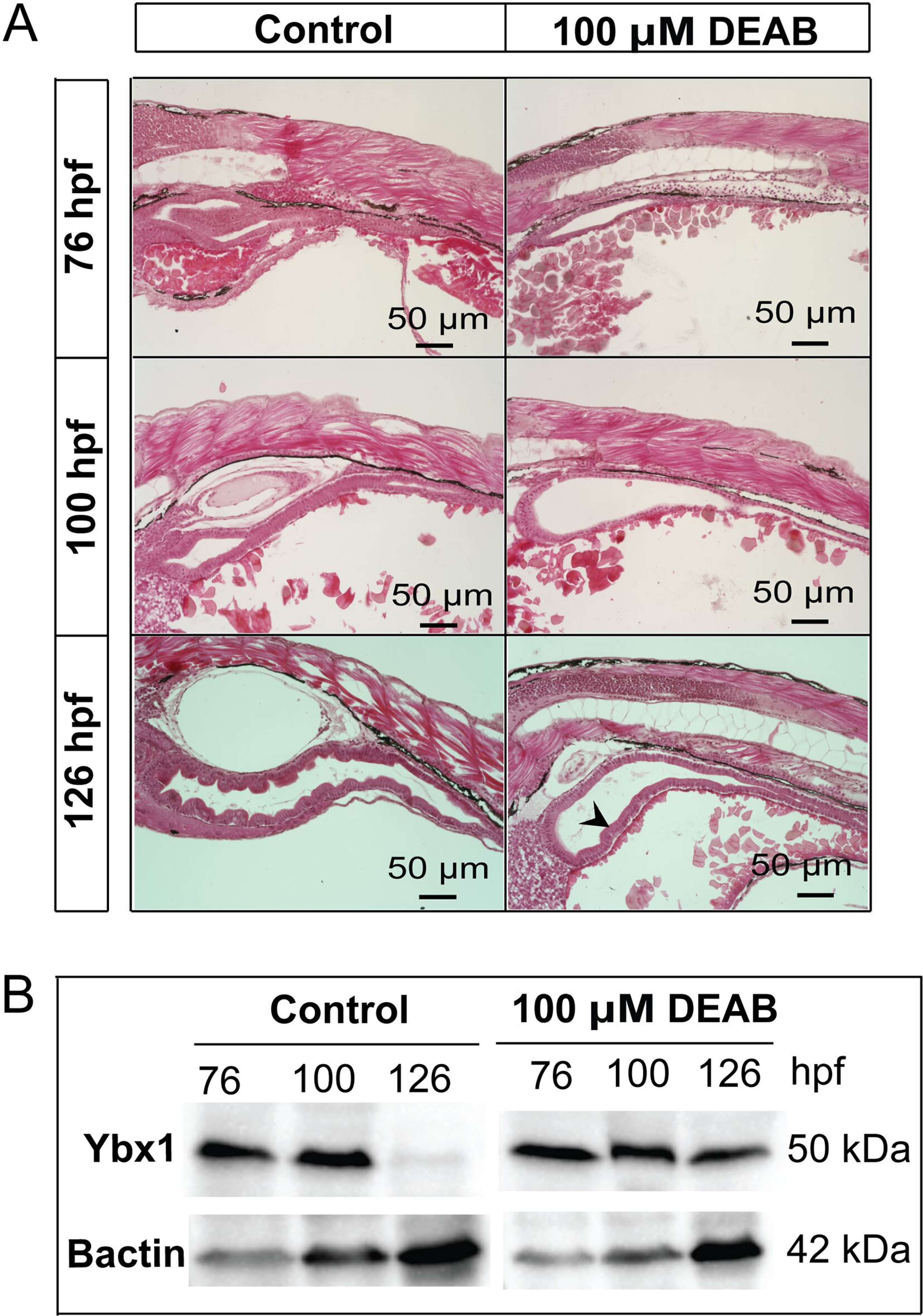
Effect of DEAB treatment on Ybx1 degradation and intestinal development. **(A)** Histological sections of zebrafish larvae treated with vehicle (Control) or 100 µM DEAB at 76, 100, and 126 hpf. Control larvae exhibited normal intestinal development by 126 hpf. In contrast, DEAB-treated larvae showed flattened intestinal surface (arrowhead). **(B)** Western blot analysis of Ybx1 (50 kDa) protein abundance in control and DEAB-treated larvae at 76, 100, and 126 hpf. Bactin (42 kDa) was used as the loading control. The data show that while Ybx1 was normally degraded by 126 hpf in controls, DEAB treatment prevented this decline, maintaining high levels of Ybx1 protein.

**Table S1.**
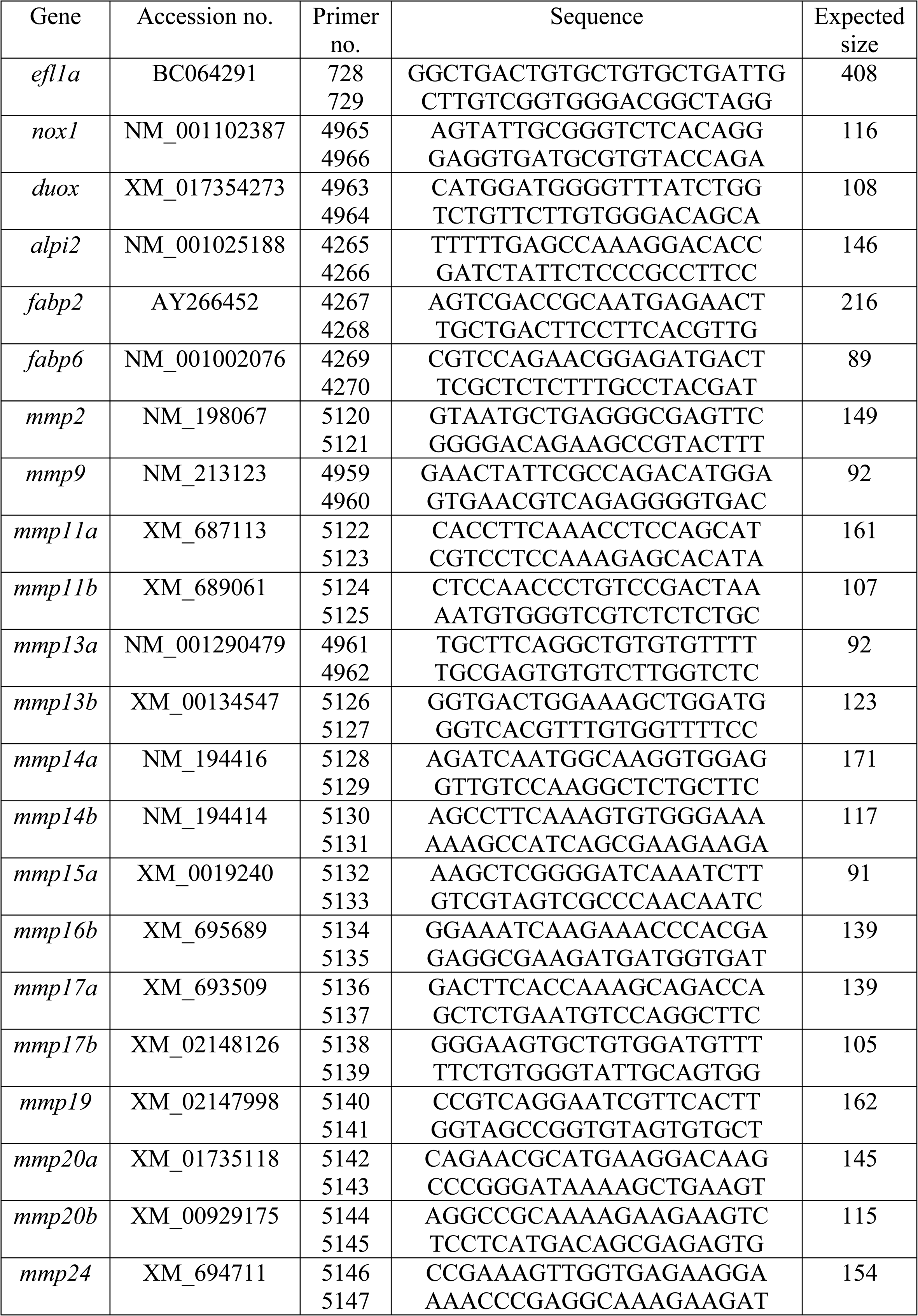
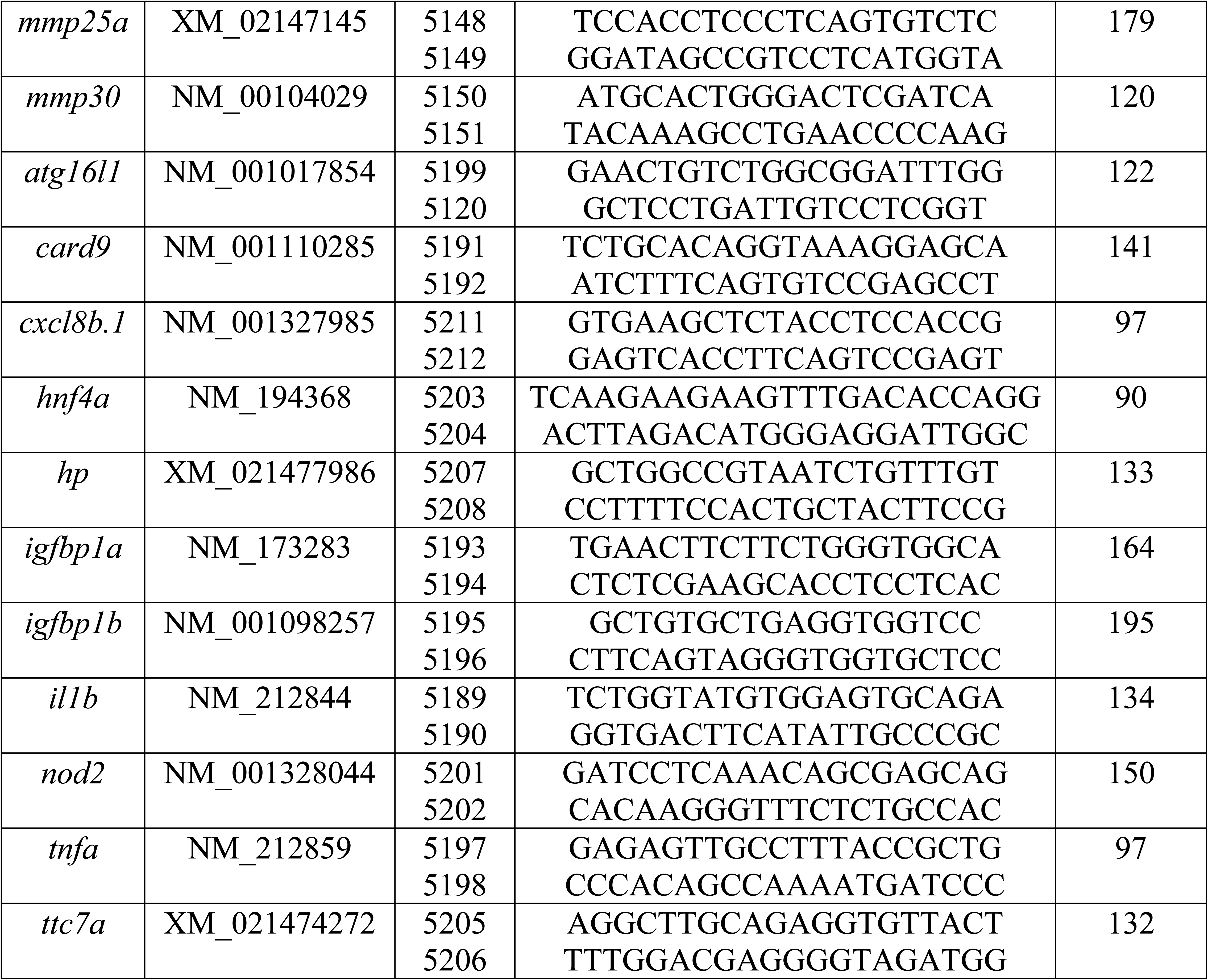
Primers used.

